# From infection to immunity: understanding the response to SARS-CoV2 through in-silico modeling

**DOI:** 10.1101/2020.12.20.423670

**Authors:** Filippo Castiglione, Debashrito Deb, Anurag P. Srivastava, Pietro Liò, Arcangelo Liso

## Abstract

**Background:** Immune system conditions of the patient is a key factor in COVID-19 infection survival. A growing number of studies have focused on immunological determinants to develop better biomarkers for therapies.

**Aim:** The dynamics of the insurgence of immunity is at the core of the both SARS-CoV-2 vaccine development and therapies. This paper addresses a fundamental question in the management of the infection: can we describe the insurgence (and the span) of immunity in COVID-19? The in-silico model developed here answers this question at individual (personalized) and population levels.

We simulate the immune response to SARS-CoV-2 and analyze the impact of infecting viral load, affinity to the ACE2 receptor and age in the artificially infected population on the course of the disease.

**Methods:** We use a stochastic agent-based immune simulation platform to construct a virtual cohort of infected individuals with age-dependent varying degree of immune competence. We use a parameter setting to reproduce known inter-patient variability and general epidemiological statistics.

**Results:** We reproduce *in-silico* a number of clinical observations and we identify critical factors in the statistical evolution of the infection. In particular we evidence the importance of the humoral response over the cytotoxic response and find that the antibody titers measured after day 25 from the infection is a prognostic factor for determining the clinical outcome of the infection.

Our modeling framework uses COVID-19 infection to demonstrate the actionable effectiveness of simulating the immune response at individual and population levels. The model developed is able to explain and interpret observed patterns of infection and makes verifiable temporal predictions.

Within the limitations imposed by the simulated environment, this work proposes in a quantitative way that the great variability observed in the patient outcomes in real life can be the mere result of subtle variability in the infecting viral load and immune competence in the population.

In this work we i) show the power of model predictions, ii) identify the clinical end points that could be more suitable for computational modeling of COVID-19 immune response, iii) define the resolution and amount of data required to empower this class of models for translational medicine purposes and, iv) we exemplify how computational modeling of immune response provides an important view to discuss hypothesis and design new experiments, in particular paving the way to further investigations about the duration of vaccine-elicited immunity especially in the view of the blundering effect of immunesenescence.

## 1 Introduction

The global pandemic set up by the severe acute respiratory syndrome coronavirus 2 (SARS-CoV-2) in the early months of the year 2020 has reached considerable proportions and, to date, does not show signs of slowdown when considered globally. In fact, as of 9:45am CET, 19 December 2020, there have been 73,996,237 confirmed cases of COVID-19, including 1,663,474 deaths, reported to WHO (https://covid19.who.int).

The mortality rates, of the SARS-CoV-2 greatly differ across the globe, ranging from 0.5 to 13% (Johns Hopkins University, https://coronavirus.jhu.edu/data/mortality), as a result of many factors including the ability to react to the pandemic by the various national health systems.

The COVID-19 disease has a quite variable clinical presentation: while the majority of individuals present with a very mild disease, often asymptomatic, a few patients develop a life-threatening disease requiring intensive care. The strongest determinant of disease severity is age, with children presenting almost exclusively with mild disease, while individuals over 70 years of age are much more likely to develop severe COVID-19 (Reference?). This variation is likely due to both host and pathogen factors. Host factors may include differences in the immune response due to genetic determinants and immunological history. On the other hand, pathogen factors include transmission, entry and spread within the host, cell tropism, virus virulence and consequent disease mechanisms.

To better understand what impact these factors may have in the differences observed in the host response to SARS-CoV-2, we set up the analysis of the dynamics generated by a computer model that considers both, the magnitude of the viral harm, and the subsequent innate and adaptive response set up to attempt achieving control of the infection. Thus we used computer simulations to create a virtual cohort of infected individuals to study the effects on the pathogenesis of both host and pathogen factors. Note that this approach goes beyond the machine learning paradigm as the knowledge is generated through a set of equations/algorithms confirmed by the scientific literature and by past models. The simulation allows systems-level, multi evidence analyses to simultaneously capture the dynamics of the major immune cell populations and the many protein mediators by which cells communicate, in order to sort out the determinants of disease severity.

## 2 Simulating SARS-CoV-2 course of events in the host

The simulation model that we used in this study, C-IMMSIM, is a derivation of an established software system for the simulation of the immune response to generic pathogens [1]. It is equipped with a fundamental innate immune response consisting in macrophages, dendritic cells and natural killer cells, and represents the adaptive immunity by B lymphocytes, plasma B antibody-producing lymphocytes, CD4 T helper and CD8 cytotoxic T lymphocytes. It is a polyclonal model as it embodies the primary sequences of binding sites of B-cell receptors (BCRs) and T-cell receptors (TCR), as well as the peptides and epitopes of the infectious agent (i.e., the SARS-CoV-2 in this case).

It represents a portion of, i) primary lymphatic organs where lymphocytes are formed and mature (i.e., mainly the red bone marrow and the thymus gland), ii) secondary lymphoid organs (e.g., a lymph node), which filters lymph, and where naïve B and T-cells are presented to antigens, and, iii) peripheral tissue which is dependent on the pathogen considered (in this case the lung).

While primary organs are just the source of lymphocytes equipped with a randomly generated receptor (actually only its complementarity-determining region, CDR), the secondary organs and the tissue are mapped onto a three-dimensional Cartesian lattice. The thymus is implicitly represented by the positive and negative selection of immature thymocytes [2] before they enter the lymphatic system [3], while the bone marrow generates already immunocompetent B lymphocytes.

C-IMMSIM incorporates several working assumptions or immunological theories, most of which are regarded as established mechanisms, including the clonal selection theory of Burnet [4], [5], the clonal deletion theory (e.g., thymus education of T lymphocytes) [2], [6], the hypermutation of antibodies [7]–[9], the replicative senescence of T-cells, or the Hayflick limit (i.e., a limit in the number of cell divisions) [10], [11], T-cell anergy [12], [13] and Ag-dose induced tolerance in B-cells [14], [15], the danger theory [16]–[18], the (generally unused) idiotypic network theory [19], [20].

Being more a general-purpose modeling platform rather than just a model, C-IMMSIM lend itself to characterise the role of the immune response in different human pathologies such as viral infections (e.g., HIV or EBV [21], [22]), cancer immuno-prevention [23], [24], type 1 hypersensitivity [25] and the chronic inflammation associated to type 2 diabetes [26] as well as specific aspects of the immune dynamics such as lymphocytes homing in lymph nodes [27], the gene regulation leading to cell differentiation [28], clonal dominance in heterologous immune responses [29] and also vaccination-eliciting fish immunity [30]. Relevant to the present study, the model has recently being used to simulate the response to a multi-epitope vaccine against SARS-CoV-2 [31], [32].

In C-IMMSIM each simulated time step corresponds to eight hours of real life. Cells diffuse randomly in the represented volume and interact among them. Upon specific recognition through receptor bindings, they perform actions which determine their functional behaviour. These actions are coded as probabilistic rules and define the transition of the interacting cell entities from one “condition” to another. In fact, each rule is executed only if the parts involved are in specific states (e.g., naïve, active, resting, antigen-presenting).

Besides cell-cell interaction and cooperation, this model simulates the intra-cellular processes of antigen uptake and presentation. Endogenous antigens are fragmented and combined with MHC class I molecules for presentation on the cell surface to CTLs’ receptors (this is the cytosolic pathway), whereas exogenous antigens are degraded into small pieces, which are then bound to MHC class II molecules for presentation to T helpers’ receptors (this is the endocytic pathway).

The stochastic execution of the algorithms coding for the dynamical rules, results in a sequence of cause/effect events culminating in the production of effector immune cells and setup of immunological memory. The starting point of this series of events is the injection of an antigen which, here, consists of the virus. This may take place any time after the simulation starts (the sequence of events of the SARS-CoV-2 simulation is reported in Appendix A). Initially the system is “naïve” in the sense that there are neither T and B memory cells nor plasma cells and antibodies. Moreover, the system is designed to maintain a steady state of the global population of cells (homeostasis), if no infection take place.

Besides the parameters defining the characteristics of the virus related to attachment, penetration, replication and assembly (i.e., its fitness), the SARS-CoV-2 virus in this model is defined as a set of B-cell epitopes and T-cell peptides consisting of amino acid 9-mers and defining its antigenicity. If the infection is stopped or becomes persistent or even kills the virtual patient it depends on the dose of the virus, its fitness, and the strength of the immune response aroused. All of these variables determine if, and to what degree, the success of the immune system requires the cooperation of both the cellular and the humoral branch, as shown in past simulation studies [33].

With respect to previous version of the model described in [34] the most important difference is that to improve the peptide-prediction performance, rather than using position-specific-score-matrices (PSSM) to weight the binding contribution of the amino-acids composing the protein segments in the bounds [35], [36], we resort to pre-computed ranked lists of T-cell epitopes calculated with the original neural network NetMHCpan method [37]–[39]. This feature, which is described below, follows the choice of a definite HLA set.

### 2.1 Selecting the HLA haplotype

The C-IMMSIM model accounts for differences in the HLA haplotype when determining which peptides are presented by antigen presenting cells. To this end, it takes in input a list of such peptides for each HLA molecule considered together with a propensity of each peptide to bind to it. This list is computed by using third party immunoinformatics tools as described in the next section 2.2.

The “HLA haplotype freq search” in the “Allele Frequency Net Database” (http://www.allelefrequencies.net) was used in order to select two HLA-A, two HLA-B and two DRB alleles which are most prevalent in US population [40]. The result pointed to the following alleles: HLA-A*02:01, HLA-A*24:02, HLA-B*35:01, HLA-B*40:02, DRB1*07:01 and DRB1*15:01.

### 2.2 Computing the peptide immunogenicity

The strain of SARS-CoV-2 used in this study corresponds to the reference sequence *NCBI Reference Sequence: NC_045512*.*2*. The primary structure of these proteins have been used to identify cytotoxic T peptides (CTL peptides) and helper T peptides (HTL peptides). To this aim we have employed two immunoinformatics tools. In particular, for the definition of CTL epitopes, the “ANN 4.0 prediction method” in the online tool `MHC-I binding prediction` (http://tools.iedb.org/analyze/html/mhc_binding.html) of the *IEDB Analysis Resource* was used for the prediction of 9-mer long CTL peptides which had an affinity for the chosen set of HLA class I alleles (i.e., HLA-A*02:01, HLA-A*24:02, HLA-B*35:01 and HLA-B*40:02) [41]–[43]. The peptides were classified as strong, moderate and weak binders based on the peptide percentile rank and IC50 value. Peptides with IC50 values <50 nM were considered to have high affinity, <500 nM intermediate affinity and <5000 nM low affinity towards a particular HLA allele. Also, lower the percentile rank, greater is meant the affinity [41]– [43]. The list of peptides is reported in the Appendix B.

For what concerns the HTL peptides, the `NetMHCIIpan` 3.2 server (www.cbs.dtu.dk/services/NetMHCIIpan) was used for the prediction of 9-mer long HTL peptides which had an affinity for the HLA class II alleles (i.e., DRB1*07:01 and DRB1*15:01) used in this study [44]. The predicted peptides were classified as strong, intermediate and non-binders based on the concept of percentile rank as given by NetMHCII pan 3.2 server with a threshold value set at 2, 10 and >10% , respectively. In other words, peptides with percentile rank ≤2 were considered as strong binders whereas a percentile rank between 2 and 10% designate moderate binders; peptides with percentile score >10 are considered to be non-binders [44]. The list of CTL and HTL peptides and the relative affinity score is reported in the Appendix C.

### 2.3 Quantifying the immunological competence

It is widely accepted that ageing is accompanied by remodelling of the immune system. With time, there is a decline in overall immune efficacy, which manifests itself as an increased vulnerability to infectious diseases, a diminished responses to antigens (including vaccines), and a susceptibility to inflammatory diseases. The most important age-associated immune alteration is the reduction in the number of peripheral blood naïve cells, accompanied by a relative increase in the frequency of memory cells. These two alterations, are extensively reported in the literature and account for the immune repertoire reduction [45], [46]. Along with the process called “inflamm-aging”, the reduction of immune repertoire is considered the hallmark of immunesenescence [47].

To model the reduction of immune efficacy we first defined the parameter “immunological competence” *IC*∈(0,1] and assumed it in a simple linear relationship with the age. Specifically we set *IC* ≡ *IC*(*age*) = −*α* · *age* + 1 with the value of the parameter *α* = 45 · 10^−4^ determined using epidemiological data as described below. Given the age, the parameter *IC* is then used to modulate both innate and adaptive immunity as follows: i) the phagocytic activity of macrophages and dendritic cells, represented by a probability to capture a viral particle, is rescaled respectively as *p*_*M*_ = *IC* · *u* and *p*_*DC*_ = *IC* · *v*, where *u*∼*U*_[*a,b*]_ and *v*∼*U*_[5×*a*,5×*b*]_ are two random variables uniformly distributed in the ranges [*a, b*] and [5 · *a*, 5 · *b*] with *a* = 25 × 10^−4^ and *b* = 10^−2^; ii) as for the adaptive immunity it is adjusted according to the immunological competence parameter *IC* by decreasing the lymphocyte counts (hence B, Th and Tc) to reflect a reduction in the repertoire of “naïve” cells with immunological history due to accumulation of memory cells filling the immunological compartment. In particular, the number of white blood cells *N* is computed as *N*∼*IC* · 𝒩(*μ*, σ^-^) = 𝒩(*IC · μ, IC*^2^ σ^2^) where 𝒩(*μ*, σ^-^) is a normal distribution with average *μ* and standard deviation σ (for each lymphocyte type B, Th and Tc) chosen to reflect the reference leukocyte formula for an average healthy human adult (see Figure 2) [48].

**Figure 1.**
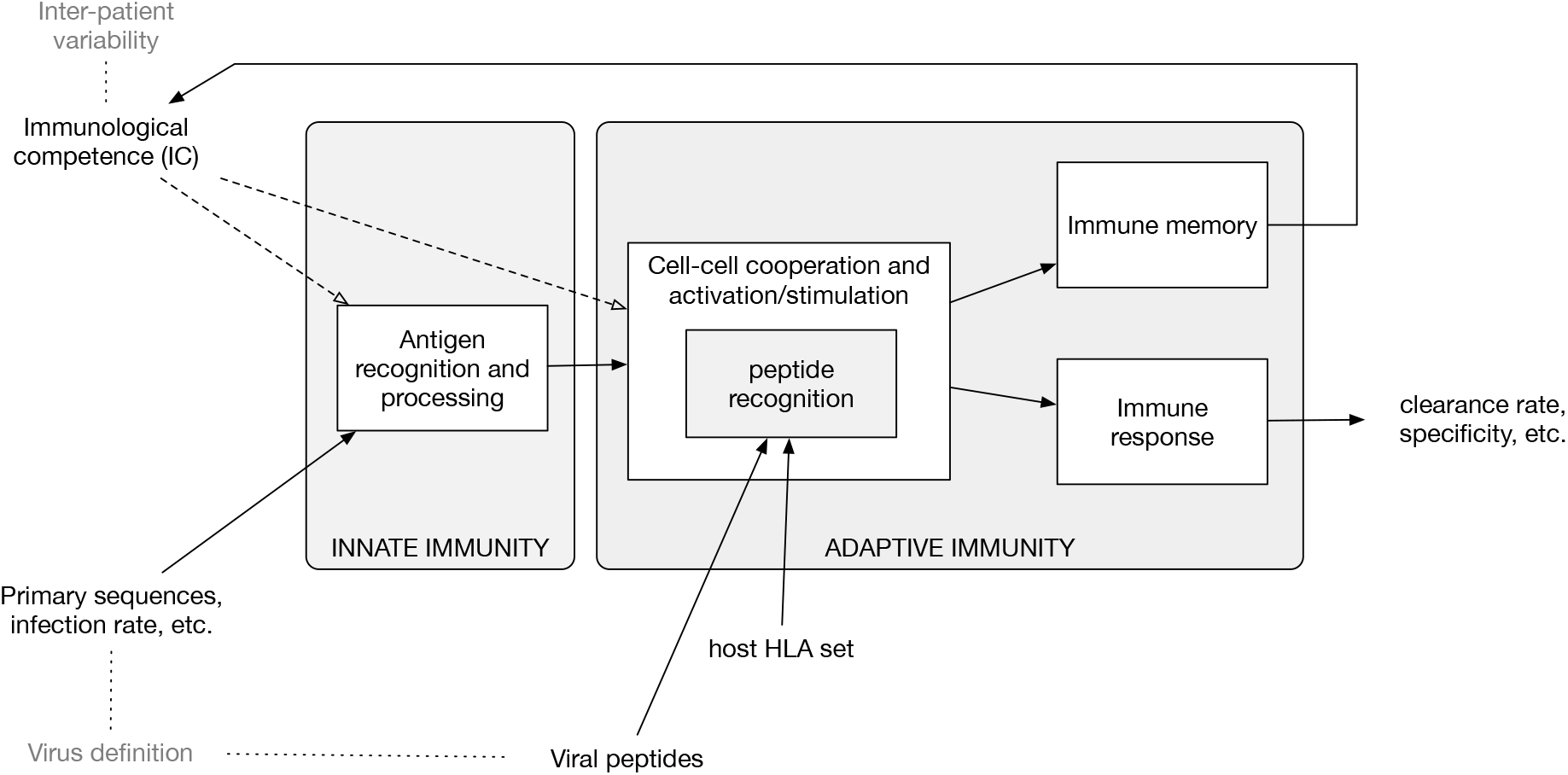
Diagram of the *in-silico* model components and accepted input. The model embodies functions to calculate the clonal affinity to precomputed viral peptides of the selected pathogen (defined by its primary sequence) with respect to a specific HLA set. The population-dynamics of the elicited lymphocytes clones, resulting from the infection by the SARS-CoV-2, provides a varying degree of efficiency of the immune response which, as it turns out, correlates with the parameters defining both the immunological competence (IC) of the virtual host and the virus definition.

**Figure 2.**
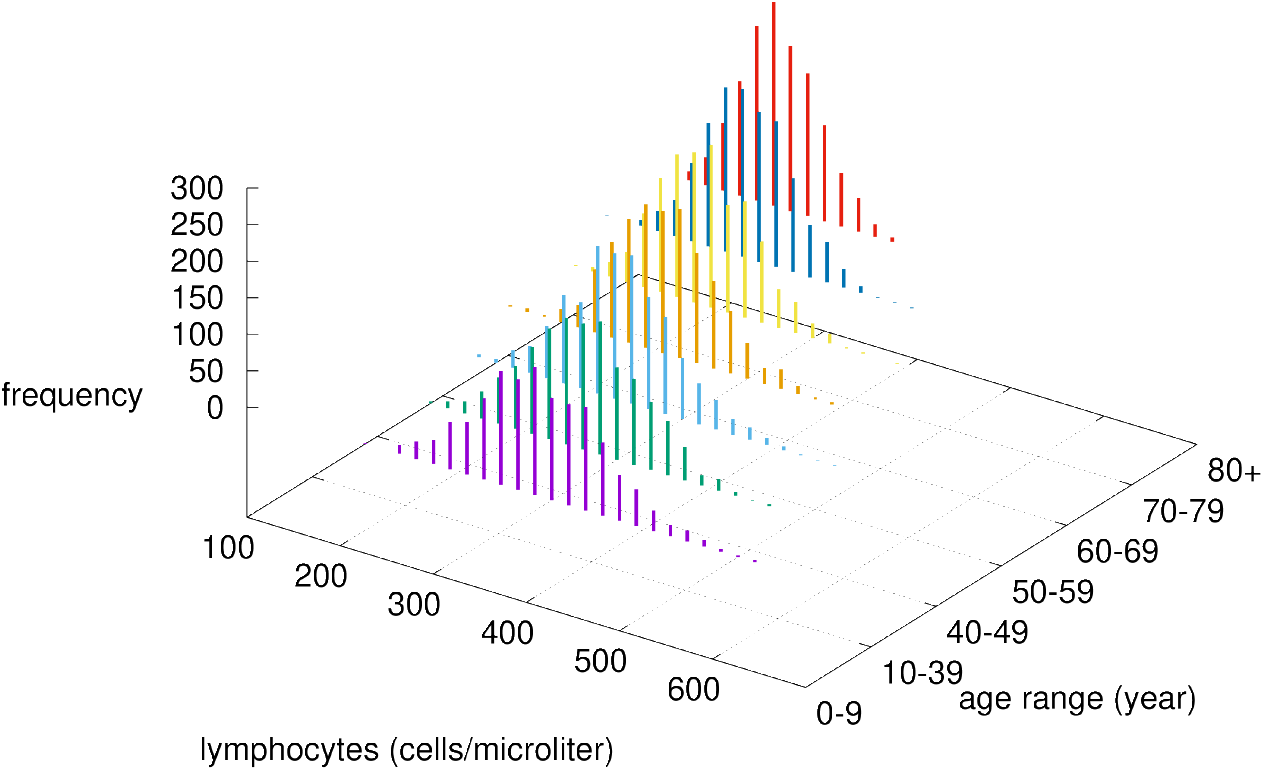
Modified lymphocytes counts by age class. Given the chosen value of *α*, the immunological competence is less than one, therefore by increasing the age, the cell numbers are drawn by a normal distribution with reduced mean value *μ* · *IC*(*age*) and reduced variance (σ · *IC*(*age*))^2^.

### 2.4 Adapting the model to SARS-CoV-2 characteristics

The infection and the dynamical features of the SARS-CoV-2 viral strain has been characterised by two parameters: i) *V*_0_ corresponding to the infectious viral load at time zero, and, ii) the affinity of the virus spike protein to the ACE2 receptor on target cells, called *p*_*A*_. In particular, 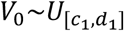 has been taken randomly in the interval [*c*_1_ = 5, *d*_1_ = 5 · 10^5^], while 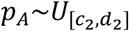 in the interval [*c*_2_ = 10^−3^, *d*_2_ = 10^−1^].

Upon choice of the age class determining the immunocompetence value *IC*(*age*) hence *p*_*A*_ and *p*_*DC*_ as well as the lymphocyte counts, the simulations depict the immune-virus competition eventually culminating in a successful, or not, virus-clearing response controlling its growth. Sometimes this control is not perfectly efficient. In those cases the result is a longer viral persistence possibly going much beyond the length of the observation period of 30 days (cf. Figure 4).

The sequence of events from viral infection leading to a full fledged immune response is detailed in the Appendix A. At each time step of the simulation C-IMMSIM dumps all variables allowing for a detailed analysis of the dynamics. A full output example of a simulation is reported and described in the Appendix D.

## 3 Modeling a representative cohort of infected individuals

We have simulated a large number of infections by varying the parameters identifying both the viral characteristics and the individual immunological competences. The seven age classes considered were 0-9, 10-39, 40-49, 50-59, 60-69, 70-79 and 80+. As already mentioned, the age class determines the immunocompetence parameter *IC* which, in turn, sets *p*_*M*_ and *p*_*DC*_ as well as the lymphocytes counts in the *in-silico* individual, we can characterize each simulation by the set of parameters (*IC*(*age*), *V*_0_, *p*_*A*_). Moreover, due to the stochasticity of the model depending on the random number realisations, each simulation corresponds to a different trajectory in the space of the variables. It follows that each simulation coincides with an *in-silico* patient with variable immunological characteristics (*IC*) and, at the same time, infected by a slightly different viral burden (*V*_0_ and *p*_*A*_).

The intervals within which these parameters vary have been chosen to reproduce the age-class incidence of disease severity of infected individuals. The age-class incidence varies wildly among regions mostly due to a different definition of COVID-19 related deaths. Moreover, due to the lack of confirmation of the causes of death in many cases in periods of high emergency as that of March/April 2020 in Italy, these rates should not be considered strictly but rather indicative of the negative exponential-like relationship of the death incidence with age. Reference values we used were the fatality rates from the the Chinese Center for Disease Control and Prevention (CDC) as of 17th February, the Spanish Ministry of Health as of 24th March, the Korea Centers for Disease Control and Prevention (KCDC) as of 24th March, and the Italian National Institute of Health, as presented in the paper by Onder et al. (2020) as of 17th March [49], [50].

To reproduce this age-related incidence of *in-silico* cases we linked the simulated viral load at a certain time to the clinical status (*clinical endpoint*). This has been done according to the rationale that a patient whose viral load is still quite high after thirty days from infection can be considered at very high risk of death. In fact in most mild cases, the clinical signs and symptoms (mostly fever and cough) have been reported to resolve within 3 weeks from the diagnosis (which translates in approximately 30 days from infection). Instead after 3 weeks several authors have described severe cases with progressive deteriorating multi-organ dysfunction with severe acute respiratory dyspnea syndrome, refractory shock, anuric acute kidney injury, coagulopathy, thrombocytopenia, and death [51].

### 3.1 Stratifying the *in-silico* cohort of patients

The analogy of some simulation variable to a realistic *clinical endpoint* allow us to stratify the in-silico patients for a more concrete interpretation of the results. We were able to classify the virtual patients on the basis of the viral load observed at day thirty (indicated by *V*_30_) and a threshold *θ* in one of the three classes:

- <u>Critical</u>: if *V*_30_ > *θ*, namely, the viral load at day 30 is still high; this class includes weak and late responders;
- <u>Partially recovered</u>: those who are still positive but have a low viral load, meaning that the immune response is controlling the viral replication (i.e., 0 < *V*_30_ ≤ *θ*); note that this class includes the asymptomatic;
- <u>Fully recovered</u> (or just Recovered): those who have cleared the virus (i.e., *V*_30_ = 0).

According to this definition by choosing the cutoff *θ* =120 viral particles per micro-litre of simulated volume, we obtain the stratification of the virtual individuals shown in panel A of Figure 3. Altogether, i.e., of all *in-silico* individuals, we get 4.3% of critical cases (broken down in age-classes in panel A), 46.8% of partially recovered (panel B), and 48.8% recovered cases (panel C). These figures sound very much in line with current epidemiological statistics when considering that the recovered cases here simulated include asymptomatics [50].

**Figure 3.**
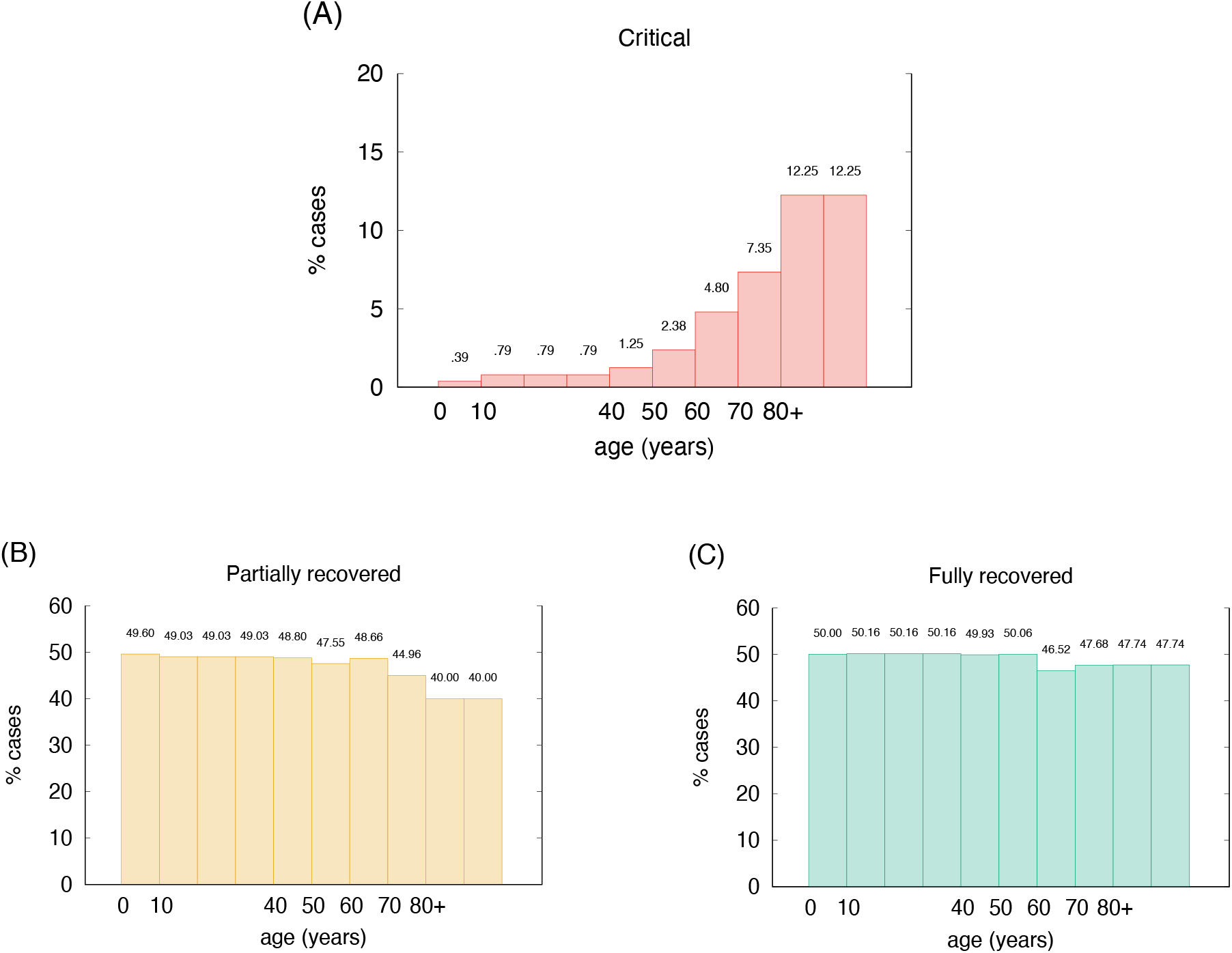
Age incidence of stratified in-silico patients (*θ* = 120 virions per micro liter). Percentages of critical cases are in agreement with [50].

**Figure 4.**
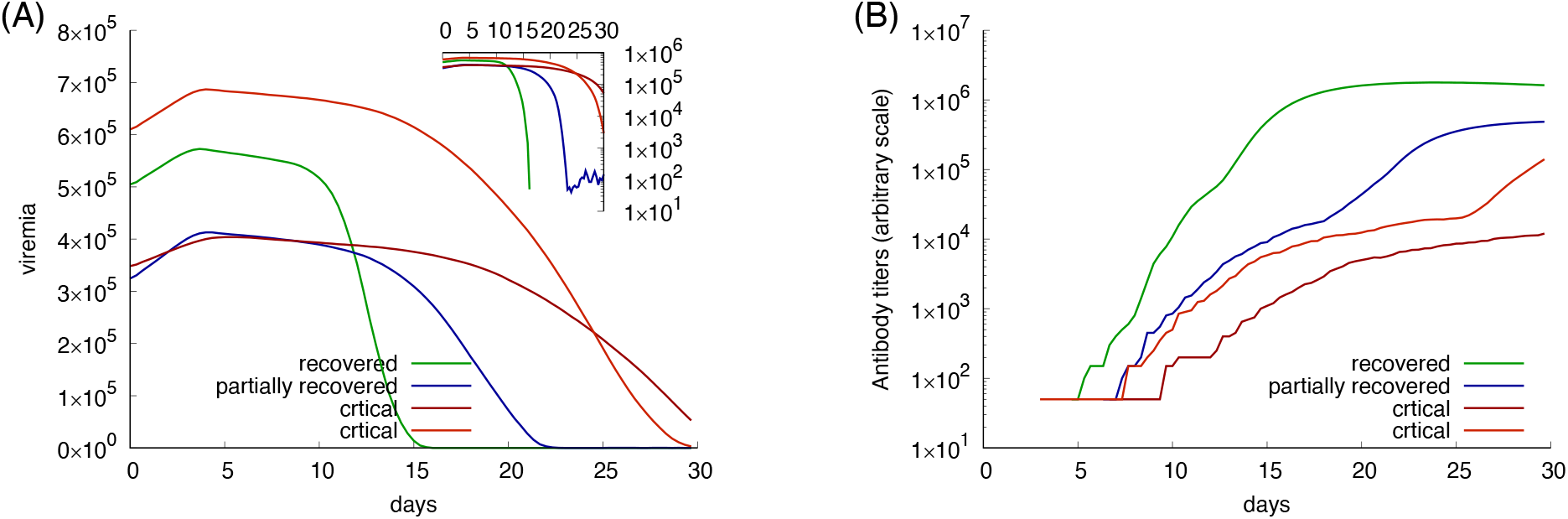
Examples of four *in-silico* cases with different outcome. Panel A shows the viral load while panel B the corresponding antibody titers. Red lines show critical cases; blue a partially recovered case; green a fully recovered case.

This results shows an interesting and surprisingly high fraction of in-silico patients in the partially recovered class. This class, in fact, includes patients that, at the end of the simulated period of thirty days, are still positive albeit manifesting an active immune response, regardless being asymptomatic or not. This question is discussed below.

These special cases can be better examined in Figure 4, which shows four distinct exemplifying runs with different outcomes. In the panel A the viremia is shown as a function of time. Red lines correspond to individuals who reach the critical condition *V*_30_ > *θ* thus falling in the class critical. The green line corresponds to a viral clearance corresponding to a fully recovered case, and the blue line shows a situation in which the virus is not completely cleared but stays below the threshold value *θ*. This case corresponds to one of what we call partially recovered as it represents virtual individuals that produce an immune response (cf. same figure, panel B showing the corresponding antibody titers) which turns out to be insufficient to clear the virus. These “unresolved infections” include asymptomatic cases and are worth the further analysis described below.

To note that the two examples of critical outcome (red curves) originate from a quite different initial viral load. Also to note that the fully recovered (green) case starts with a viral load that is higher than one critical case, still the immune response manage to control the infection. The blue curve shows a partially recovered case which greatly decrease the viral load (inset plot of panel A) but does not clear it completely.

### 3.2 How the model explains symptoms

It is worth to clarify that the term “symptom” has no meaning in the *in-silico* framework until we specify the link between model variables and possible *clinical endpoints*. Also we should note that we have no concept of comorbodity here that would help in defining the “status” of the virtual patient. To overcome this limitation, besides the viral load at day 30, we think up the following quantities (or variables) as *clinical endpoints*: (a) the damage in the epithelial compartment, namely, percent of virus-target cells that are dead at the time of observation as surrogate marker of vascular permeability; (b) the concentration of pyrogenic cytokines as a *surrogate marker* of fever, i.e., Prostaglandins TNFa, IL-1 and IL-6 causes fever people get varying degree of severity with COVID-19.

Of these two potential surrogate marker of criticality, the first appear more appropriate. In fact, while the amount of pyrogenic cytokines (surrogate clinical endpoint b.) correlates with the severity of the disease, the most striking difference in the critical cases versus the non critical (i.e., recovered plus partially recovered) is seen when comparing the accumulated damage in the epithelial compartments (*ϕ*) computed as the fraction of depleted epithelial cells due to the immune cytotoxicity of SARS-CoV-2 infected cells during the whole observation period (surrogate clinical endpoint a.),

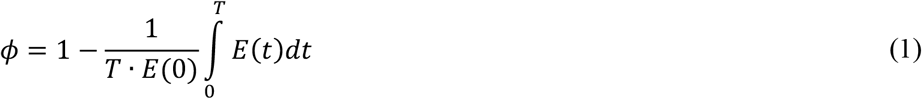

where *E*(*t*) is the epithelial count per micro litre of simulated volume and *T* is the time horizon of 30 days (note that *ϕ* ∈ [0,1]). Indeed, when we plot the distribution of *ϕ* for the cases in the critical and non critical (i.e., partially recovered plus fully recovered) classes separately we obtain what shown in Figure 5. The plot clearly shows that for critical in-silico patient the damage is much more pronounced than for non critical ones. This prompt us to use the threshold *ϕ*_*c*_ = 0.63 to set apart patients which have mild infections (about 80% as in reality [52]–[54]) to those having severe disease (15% with dyspnoea, hypoxia, lung changes on images [55]) or critical illness (5%, respiratory failure, shock, multi organi dysfunction, cytokine storm syndrome [56]), that is, we label patients with *ϕ* ≥ *ϕ*_*c*_ as symptomatic while those with *ϕ* < *ϕ*_*c*_ asymptomatic. According to this further stratification patients who are still positive (i.e., 0 ≤ *V*_3._ < *θ*) and have no symptoms (i.e., *ϕ* < *ϕ*_0_) account for about 44% of the simulations which is in line with current estimates of asymptomatic incidence (Italian Ministry of Health Report, in Italian http://www.salute.gov.it/portale/nuovocoronavirus/dettaglioNotizieNuovoCoronavirus.jsp?lingua=italiano&menu=notizie&p=dalministero&id=4998).

**Figure 5.**
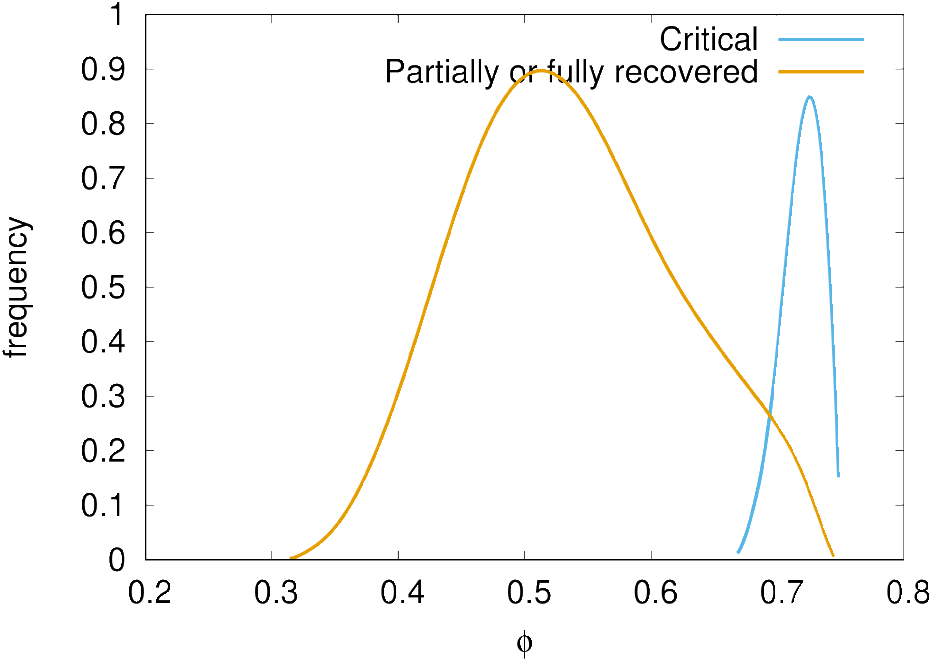
Distribution of epithelial damage *ϕ* of the cohort cases classified in critical and non-critical. A delimiting value *ϕ*_*c*_ = 0.63 separates well the two classes.

### 3.3 A high viral load carries a serious risk

We tested the correlation between the antigen abundance (or infective viral load *V*_0_) and the severity of the infection. The Mann-Whitney-Wilcoxon (MWW) test shows significant (i.e., p-value< 10^−3^) difference between infecting viral load *V*_0_ in the three classes critical, partially recovered, recovered. In particular we find that a higher *V*_0_ is a strong correlate of disease severity [57]. Of interest is the fact that there is no significant difference among age groups, that is, *V*_0_ is not predictive of the disease severity with respect to the age [58] (MWW test p-value>0.05).

### 3.4 IL-6 correlates with disease severity but youngers generate more

Significantly, in most critically ill patients, SARS-CoV-2 infection is associated with a severe clinical inflammatory picture based on a severe cytokine storm that is mainly characterised by elevated plasma concentrations of interleukin 6 [59]. In this scenario, it seems that IL-6 owns an important driving role on the cytokine storm, leading to lung damage and reduced survival [60].

The simulation agrees on this finding as the plot in Figure 6 shows. Plotting the peak value of the viral load (i.e., the maximum value attained in the observed period) versus the logarithm of the integral of IL-6 over the whole period (cf. eq(2) in next section 3.5), we see a positive correlation no matter the outcome (recovered, partially recovered, critical).

**Figure 6.**
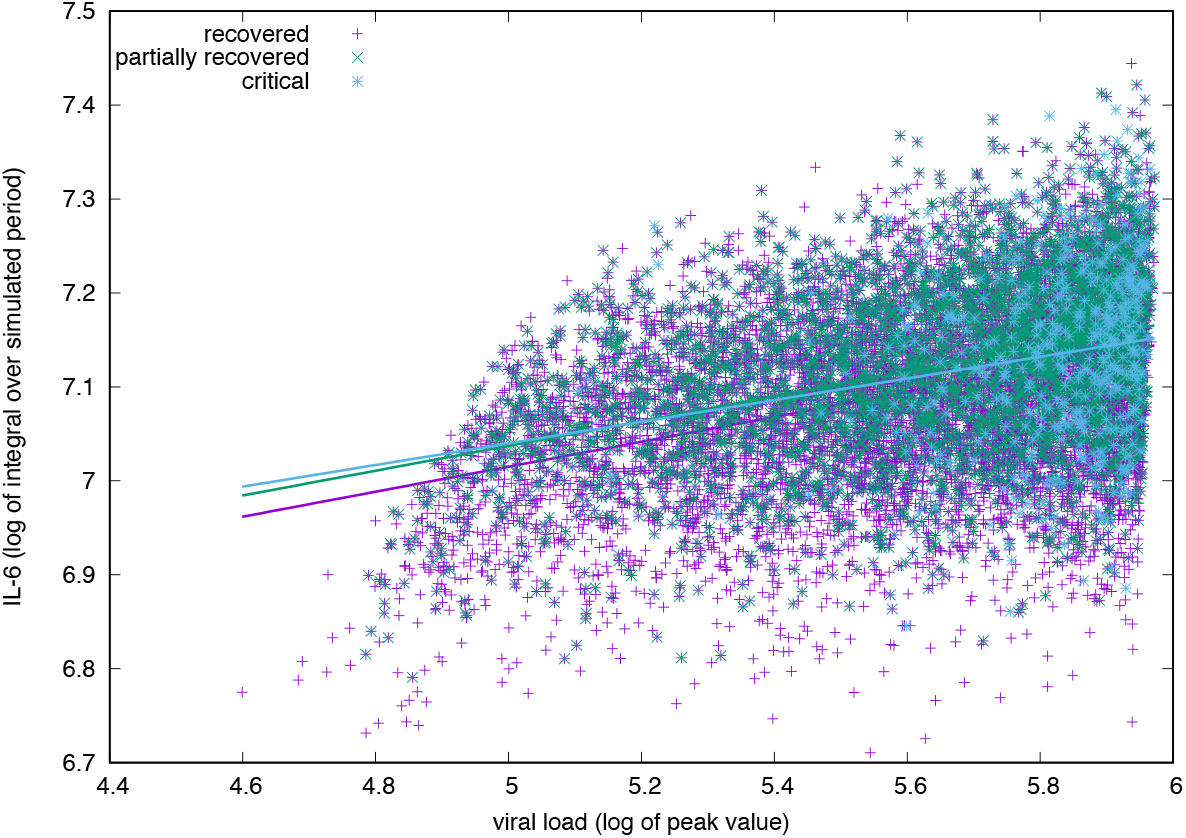
IL-6 concentration (log-scale) correlates positively with the viral load measured at the peak (i.e., its maximum value during the simulated period). The correlation is positive for all groups, critical, partially recovered and recovered with no significant difference in the degree of correlation.

For all age-classes, a critical clinical course is associated to a significantly higher concentration of pro-inflammatory cytokine IL-6. Panel B of Figure 7 shows the same information for all age-classes lumped together and the difference is statistically significant. The cytokine concentration on the y-axis is calculated as the integral over the whole simulated period (definition in eq(2) of section 3.5). The positive correlation between inflammation and severity of the clinical course is a consequence of the struggle of the immune system to cope with the infection. However what panel A of Figure 6 reports is a generic higher production of IL-6 in younger individuals compared to elders. The explanation of this outcome becomes visible following the line of consequences starting from a stronger cytotoxic activity (see panel B of Figure 12 below) that killing infected cells cause a stronger release of danger signal to which macrophages respond by secreting IL-6. Since younger have a higher immunological competence (IC), they respond with both stronger cytotoxic response and better innate (i.e., macrophage) activity. The result is the somehow counterintuitive observation that while younger individuals are more inflammed, they have a smaller propensity to experience severity of the disease.

**Figure 7.**
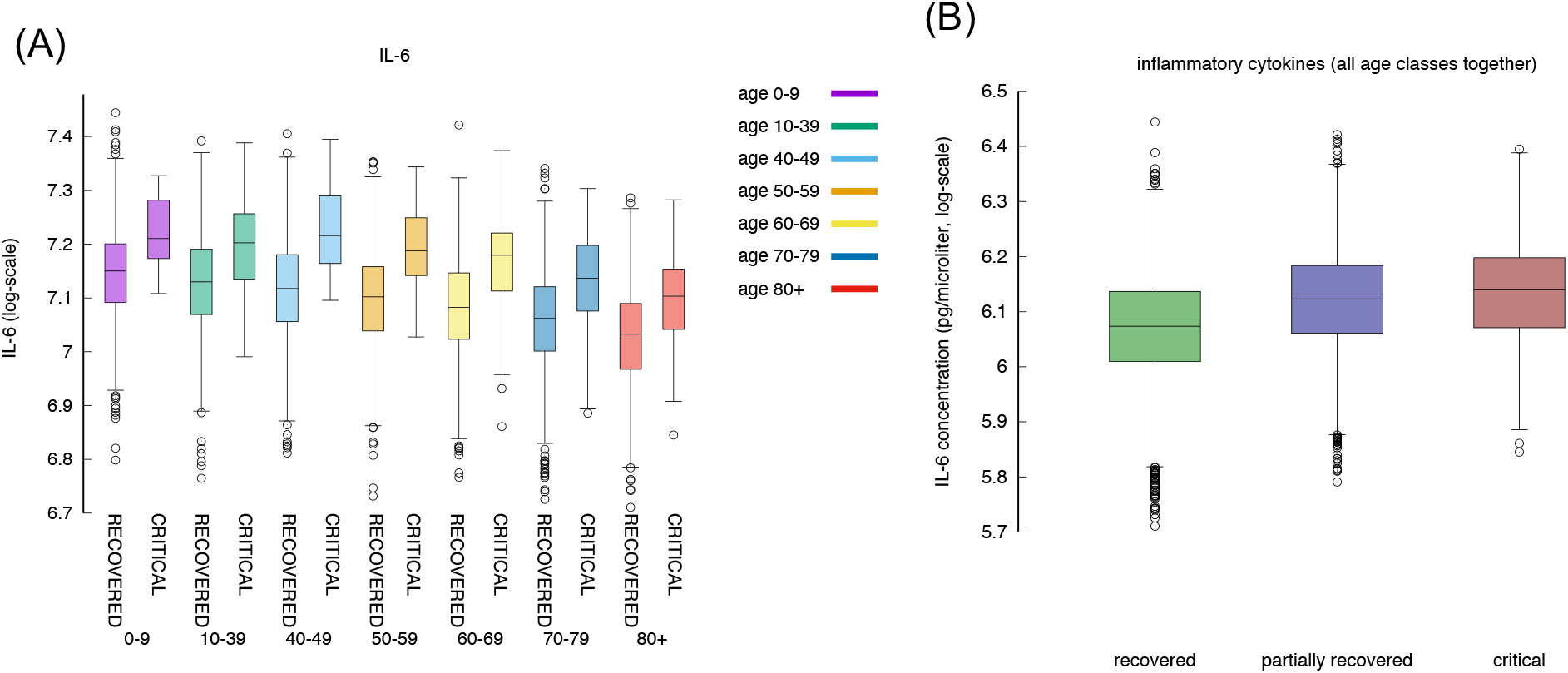
IL-6 with respect to age and severity of the disease. Inflammation correlates positively with severity of the disease (MWW test, p-value< 10^−3^) [61]. Here IL-6 is the area under the curve, as defined in section 3.5.

### 3.5 Younger individuals deal with the virus producing more cytokines

What observed in the production of IL-6 in younger individuals extends to all cytokines produced during the response to the inflammation. In fact we find that, in general, cytokines’ cumulative production during the whole simulated period correlates inversely with the age. Calling *c*_*x*_ the concentration of cytokines *x* in the simulated volume, where *x* is one of IL-6, D, IFNg and IL-12, we define

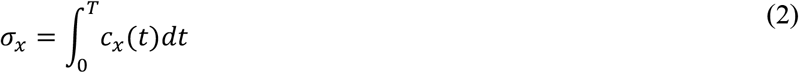

the cumulative value of cytokines in the whole observation period. Figure 8 shows σ_*IL*6_, σ_*D*_, σ_*IFNg*_, and σ_1*L*12_ with respect to age. What Figure 8 shows is that there is a clear reduction of cytokines’ production with age. This, similarly to what discussed in the previous section, is due to the reduced immune activity which indeed is determined by a reduced immunological competence with the age [62]. To justify the apparent contrast of this finding with the fact that elder acute infected individuals are more prone to experience a cytokine storm we should openly regard to one of the limitation of the model, namely, the lack of further cytokine feedbacks that are activated during the course of an extended pneumonia.

**Figure 8.**
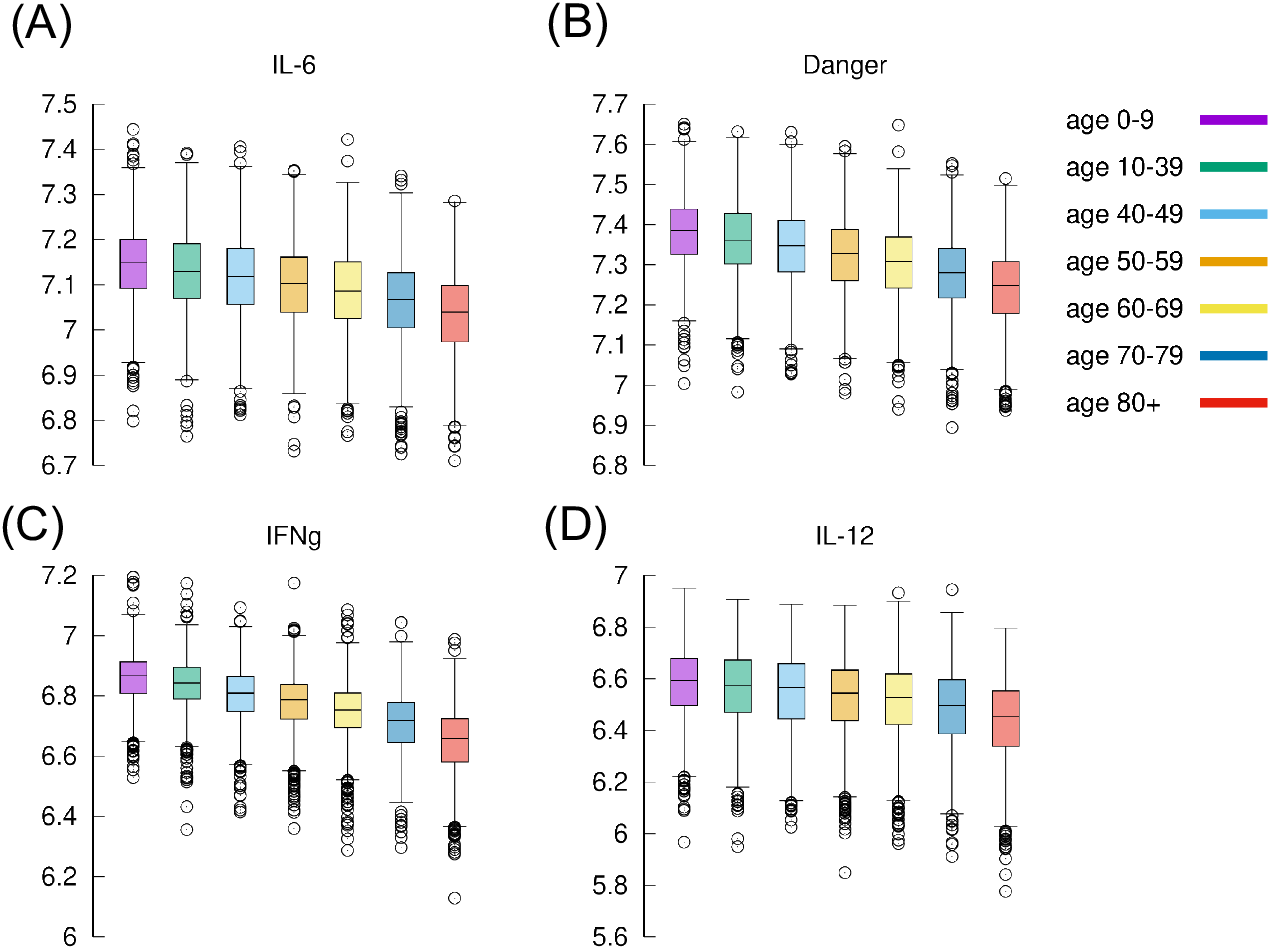
Cytokines. Shown IL-6, Danger signal, IFNg and IL-12 per each age class. All showing the same reduction with respect to increased age.

### 3.6 IFNg concentration is higher in milder courses of the infection

The expression of IFNg by CD4 tended to be lower in severe cases than in moderate cases as shown in panel A of Figure 9 and agrees with [63].

The inverse correlation of interferon-gamma (IFNg) with disease severity is observed in all age groups (panel A and also in panel C when summing all ages-classes). Interestingly, recovered and partially recvovered do not show a meaningful difference when compared to the critical cases (panel C).

**Figure 9.**
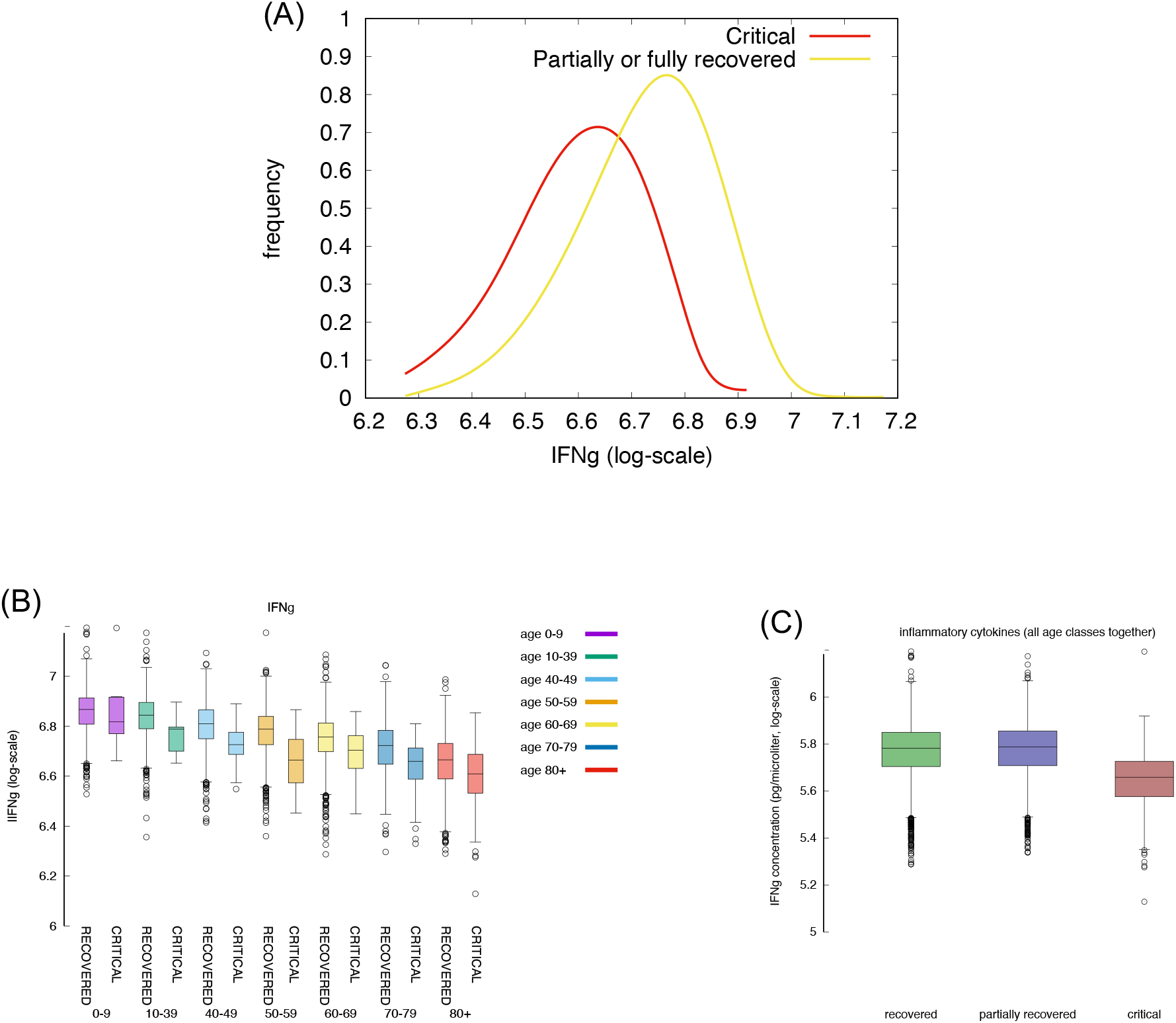
Severe cases are associated to a lower concentration of IFNg. IFNg is measured as the area under the curve (i.e., the integral in the simulation time window).

IFNg is released by natural killer (NK) cells upon bystander stimulation by danger signals (Rule n.5 in Appendix A) which, in turn, is released by infected/injured epithelial cells upon viral infection (Rule n.3) and when killed by cytotoxic cells (Rule n.18). This analysis lead us to state that a prompt activation of NK cells in younger individuals due to a higher immunological competence, and a stronger cytoxic response killing infected cells, controls the “acute” production of danger signal impacting on the production of IFNg.

### 3.7 Cytokine storm goes with symptoms

If we use the cumulative value of inflammatory cytokines as variable, namely, σ_*INFγ*_ + σ_*IL*6_ + *D* + σ_*TNFa*_ (i.e., the sum of the integrals) and we use the accumulated damage in the epithelial compartments, that is, the fraction of depleted epithelial cells due to the immune cytotoxicity *ϕ* defined in eq(1) (cf. section 3.2) as the discriminating criteria between symptomatic and asymptomatic, we observe what is shown in Figure 10.

**Figure 10.**
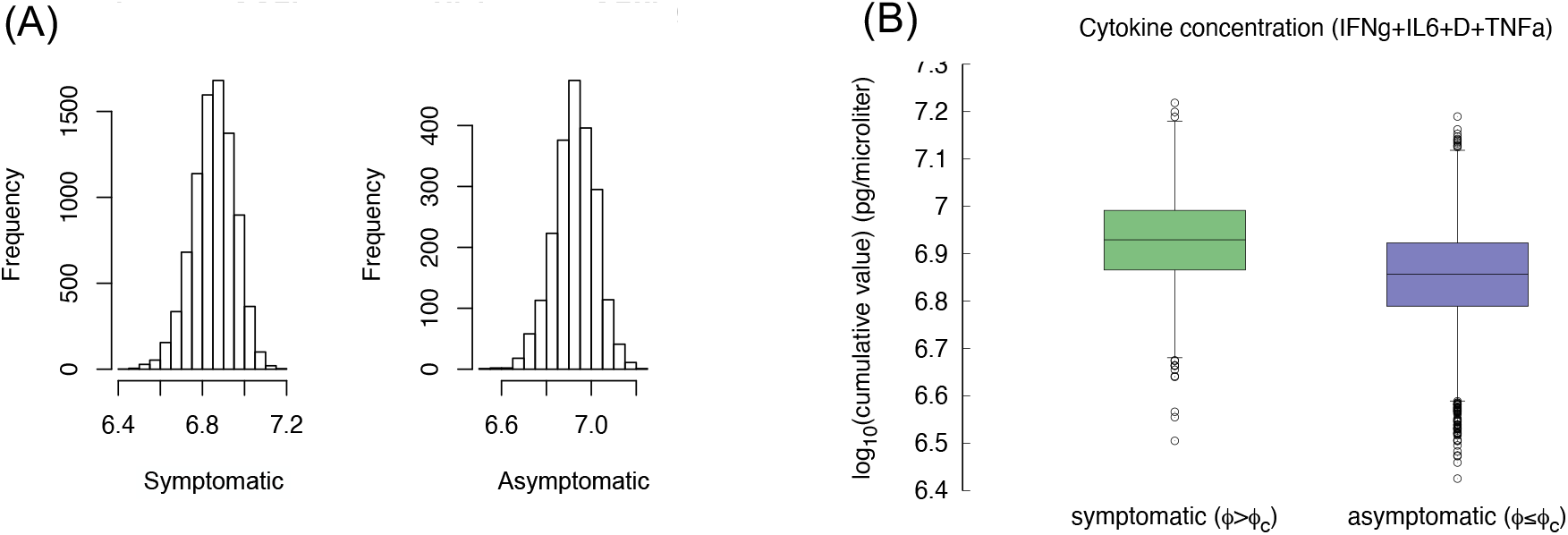
Compare the “cytokine storm” in the two groups asymptomatic *ϕ* < *ϕ*_*c*_ and symptomatic *ϕ* ≥ *ϕ*_*c*_. Panel A shows the histogram, panel B compares the wiskers. The difference is statistically significant (p-value< 10^−3^).

Compared with asymptomatic cases, the symptomatic ones more frequently have a markedly higher levels of inflammatory cytokines. The difference of the virtual patients in the two classes is statistically significant (MWW test, p-value< 10^−3^), which is in line with the clinical finding that show higher inflammatory level in severe disease progressions [63]. This result seems to contrast what stated in section 3.5, namely that younger individuals produce more cytokines but have a less-severe course of the disease. However the explanation provided by the simulation is that those who deal with the infection more rapidly (those including asymptomatics) produce, overall, a smaller amount of cytokines, thus are at lower risk of having a “cytokine storm”. On the other hand, an inconclusive immune response chronicisizing the inflammation results in pronounced symptoms (e.g., extended epithelial damage) and ultimately in a cytokine storm.

### 3.8 Why the immune response is quicker in younger individuals

It has been suggested that in younger individuals several factors contribute to the lower numbers of patients obseved with severe disease, namely: lower number of ACE receptors, overlapping immunity against coronaviruses and a more efficient intact immune system.

Indeed Figure 11 shows that the immune response is quicker in younger virtual individuals compared to elder ones. The speed of the immune response is calculated in terms of the time (in days) the viral load *V*(*t*) reaches its maximum (indicated 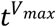 where *t*: *V*(*t*) = *V*^*max*^ and 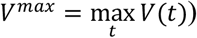 and starts to decline due to the immune response. Panel A of Figure 11 shows the distribution of 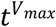 for each age class. Clearly, younger individuals develop a faster response and consequently the virus is cleared earlier. This is shown in panel B which plots the distribution of the time (in days) it takes the immune response to decrease the viral load below the threshold *θ* whenever this happen (the cases for which *V*(*t*) > *θ*, ∀*t*, are not counted in this statistics). Panel B is in line with the fact that younger individuals mount a quicker immune response that is generally more efficient than those in elder people thus eradicating the virus in a shorter time.

**Figure 11.**
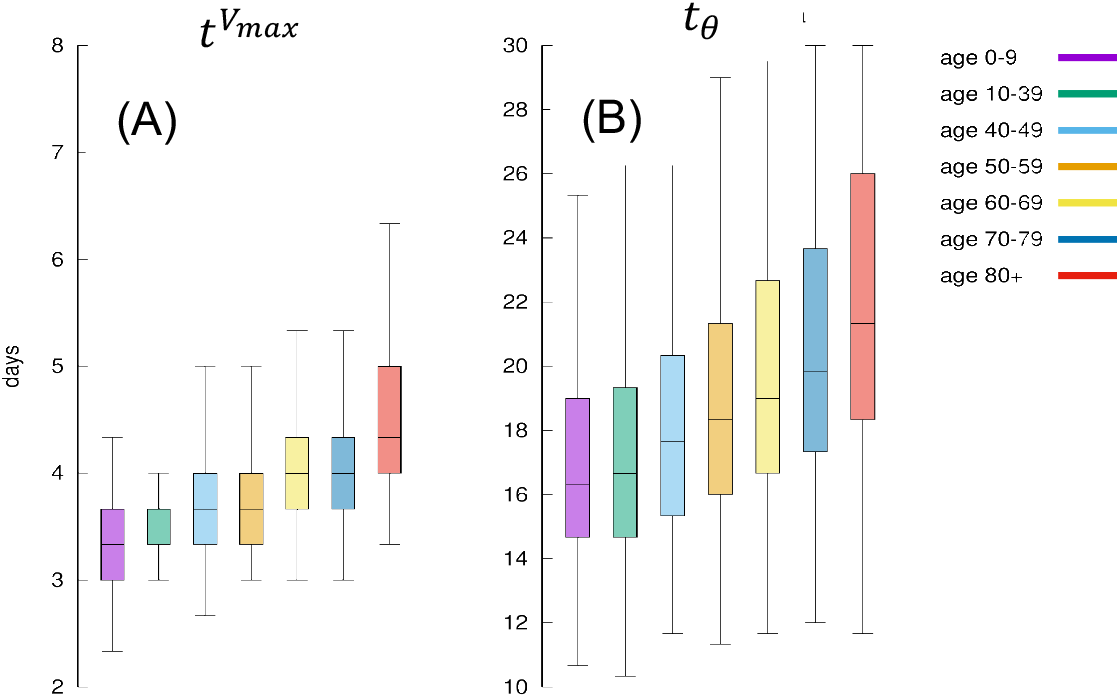
Panel A shows for each age class, the days elapsed from infection until the viral load starts to decrease. This is indicated as 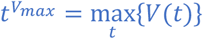 since it means the distribution of the time when the antigen reaches the peak. It is a measure of how quick the immune defences are mobilised, hence its speed. Panel B shows the corresponding distribution of 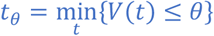 , namely the time it takes for the immune response to bring the viral load to fall below *θ*. It is a measure of the efficiency of the immune response in clearing the infection.

**Figure 12.**
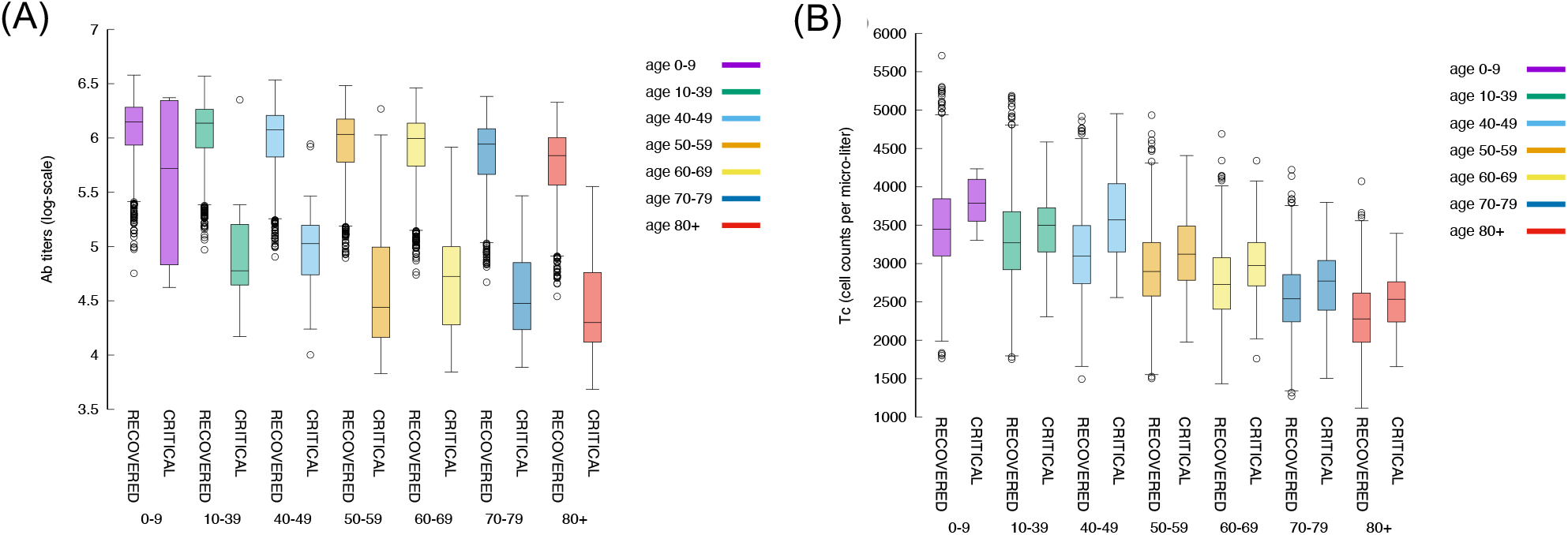
The relationship between magnitude of the immune response and age is maintained also when looking at antibodies and cytotoxic cells. Panel A shows the antibody titers for all age classes comparing recovered with critical patients. Panel B panel shows cytotoxic cell (Tc) counts (peak values).

### 3.9 The key role of the humoral response

Figure 12 shows that younger individuals have a higher production of antibodies when compared to elder individuals. This is evidente for both critical and recovered (partial or fully recovered) individuals. However the most striking observation when considering the difference between recovered and critical cases is the gap in antibody titers present in virtually all age classes (panel A). This indicates a strong protective role of the humoral response making a split between recovered and critical patients.

Panel B of Figure 12 shows the corresponding statistics for the cytotoxic T cell (peak value) count per age-class and critical status. This plot consistently evidences that youngers have a stronger response than elders. Interestingly, in contrast to the humoral response, in all age classes the cytotoxic response in critical individuals is higher than in recovered ones revealing the attempt of the immune system to counterbalance the unefficacy of the humoral response.

Moreover, a further view at the antibody titers reveals that its peak value (i.e., 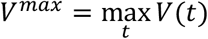 correlates inversely with the clearance time (*t*_*Ϙ*_), that is, faster response are obtained with a lower production of antibodies (cf. Figure 13). This is in line with the hypothesis that asymptomatic individuals develop a rapid but mild response which clears the infection [64].

**Figure 13.**
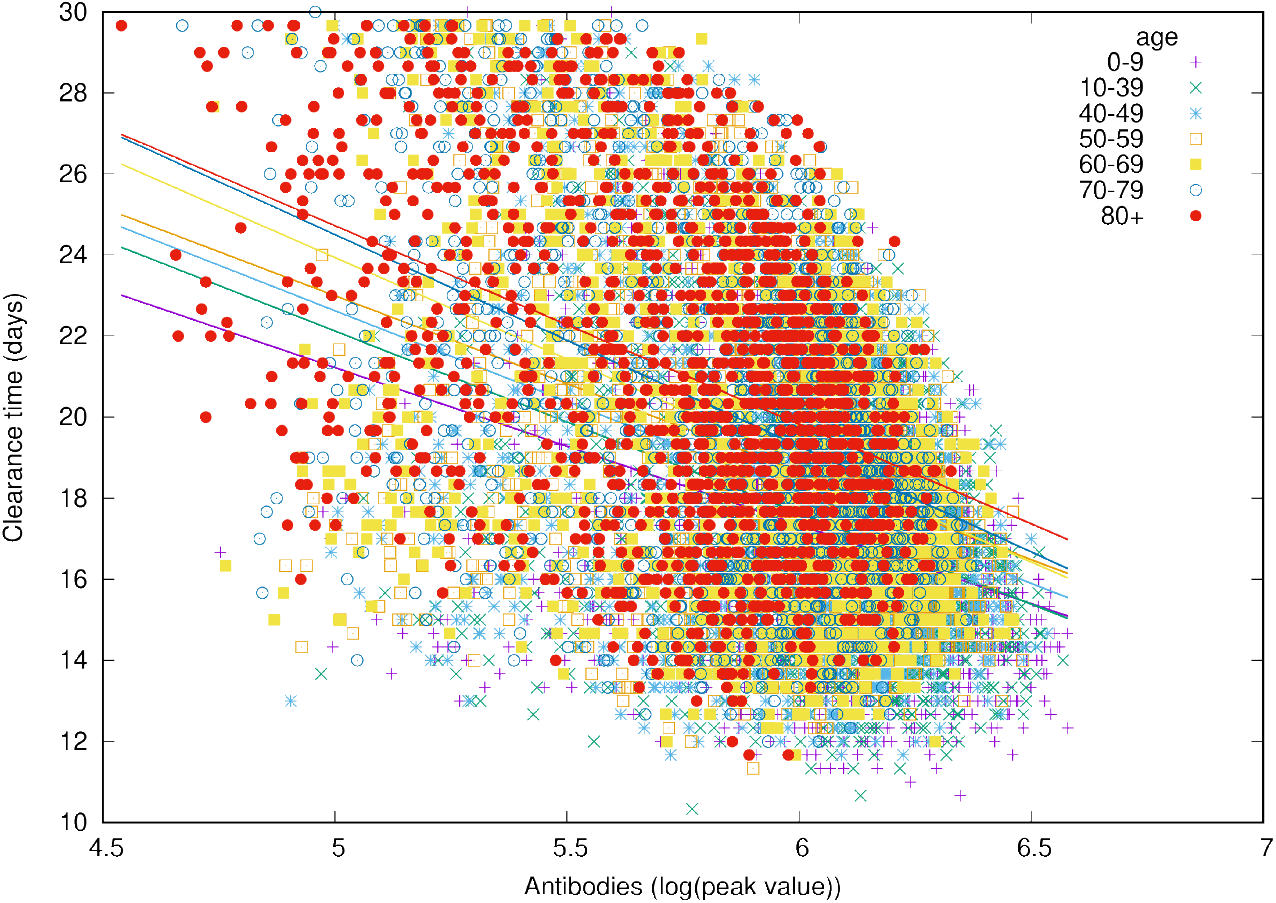
The peak value of antibody titers correlates inversely with the time-to-clear-virus *t*_*Ϙ*_.

It should be noted, however, that there are still substantial uncertainties on the data available due to the variable diagnostic accuracy of different serological tests for COVID-19, therefore more well designed large clinical studies are warranted to address this matter ([65], [66]). Interestingly, the same can not be said when cosidering peak values of Tc counts, that is, the cytotoxic response does not correlate, either positively or negatively, with time-to-clearance (not shown).

### 3.10 Antibody titers have a prognostic values after day 25

We have used a logistic regression classification to see if by measuring the antibody titers and the CTL counts at day *t* < 30 we are able to infer the outcome at the end of the simulated period of *t* = 30 days. We call *V*(*t* = 30) = *V*_30_ the viral load at day 30 after the infection.

Formally, the logistic regression classifier uses the data set {(*x*_*i*_, *y*_*i*_)}_*i*=1… *m*_ where 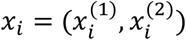 is the feature vector consisting in the normalised cytotoxic T-cell lymphocytes count, 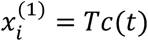 and the antibody titers at day 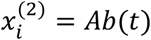. While *y*_*i*_ = 0 if the corresponding run has *V*_30_ ≤ *θ* and *y*_*i*_ = 1 if the corresponding runs has *V*_30_ > *θ*.

Panel A of Figure 14 show the features *x*_*i*_ corresponding to the recovered cases (i.e., *y*_*i*_ = 0) represented as yellow circles and the critical cases (i.e., *y*_*i*_ = 1) corresponding to black daggers. This panel shows the best separation curve found after training a logistic regression model on the training sample *x*_*i*_ = (*Tc*(25), *Ab*(25)), namely, Tc count and antibody titers measured at day 25 from infection.

**Figure 14.**
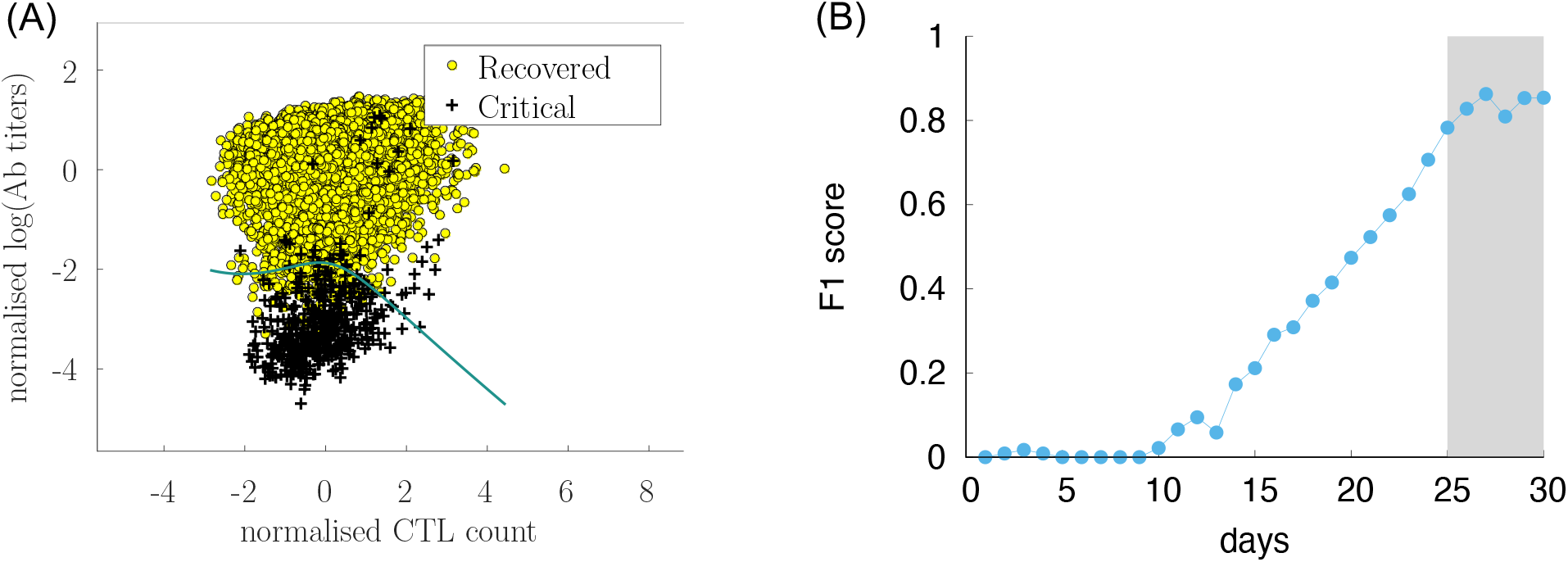
Sørensen-Dice coefficient (F1 score) of a logistic regression ML model to predict outcome (recovered/critical) from *Tc*(*t*) and *Ab*(*t*) at various days. The analysis shows that starting from day 25 from infection, the couple cytotoxic T cell counts, antibody titers is informative for predicting the outcome.

Panel A shows the data set after the classification in recovered and critical and the separation curve. In the figure the data set corresponds to the observation at day 30 while the analysis has been conducted at different time points. Panel B shows how the classification accuracy increases with the passing of time. In this panel we plot the Sørensen-Dice coefficient (most know as the F1 score [67]) which increases when the assessment is made by using features (i.e., Tc and Ab measurements) later in time as the infection and corresponding immune response develops. Interestingly, before day 10 after the infection it is not possible to find a meaningful classification criteria which predicts the outcome, while the Sørensen-Dice coefficient increases to a high value already at day 25 indicating that 25 days after infection the level of immune activation represented by the antibody titers and cytotoxic counts is predictive of the clinical outcome.

## 4 Discussion and conclusions

The immunological correlates of COVID-19 are far from being clearly elucidated in clinical studies. Simulation studies can help disentangling the importance of factors such as a reduced ability to mount an efficient (i.e., not off target) immune response due to age or the infective viral load determining the initial viral burden.

We have set up a computational model that simulates the infection with a varying dosage of the virus and with a slightly different affinity to the ACE2 receptor of target cells, in individuals with different immunological competence.

The results of a large number of simulations that we call *virtual* or *in-silico cohort*, demonstrates that the great variability observed in the real pandemic can be the mere result of such diversity in both viral and human characteristics.

The computational model used is able to explain a number of clinical observations of SARS-CoV-2 infection and to evidence the importance of the humoral response in discriminating an efficient from a poor immune response failing to completely clearing the infection and, in some cases, bringing the viral load down below a threshold value and, at the same time, without showing markers of symptoms.

The model has been tuned for parameters able to reproduce the relationship of age with the disease severity (cf. Figure 3). Starting from that, any other observation revealed an emergent property of such complex simulation environment. In particular we observe the correlations among infective viral load *V*. and severity, among immunological (in)competence (thus age) and severity, among the overall cytokine levels and symptoms (i.e., a virtual cytokine storm), and, finally, the key role of the humoral response in clearing the infection yet sustained by the cytotoxic activity (cf. Figure 12). Importantly, we have identified day 25 after infection (which we can roughly associate to about day 15^th^-18^th^ after the appearance of the symptoms) as the time for a predictive measurements of the antibody response to assess the risk of developing a severe form of the diseases. Before that time, our data suggest that the prediction is not statistically meaningful.

The model is restricted in a number of aspects. It simplifies reality and works with a limited number of mechanisms and a reduced diversity. Moreover, it does not reproduce diverse organs and tissues and therefore we cannot observe site-specific pathological problems, including the spatial extension of pneumonia. Nevertheless the analysis conducted in the present work accounts for such limitations and the results obtained can be reasonably considered independent of such restrictions.

Of course there are some cases reported in the literature in which the course of infection has been extremely long, expecially in severly immunocompromised patients. However, the very complex and lenghty dynamics taking place in those cases are beyond the scope of this study, which instead represent at large the majority of observed cases.

Finally, we should consider that the clinical ground of observation inevitably starts much later than in our model, as people ask for medical attention only after developing symptoms or after knowing of accidentally having been in contact with patients/carriers. Therefore the window of observation we consider in this paper is recapitulating more precisely the infection dynamics of the early days.

Potential study directions should cover the duration of immunity, either natural or induced by the vaccine, including ways to indirectly verify it, the impact of immunesenescence on the elicited immunity or the combined effect of monoclonal antibody and vaccines in potential future terapies.

Despite the extraordinary complexity of the immune system dynamics, the progress of simulation platforms suggests that a more intense interaction between clinicians and researchers in computational model could bring these models to the desidered quality for deployment in the medical field.

Appendices

## Appendix A. The sequence of events from virus infection to immune response

The system is in a stable state (apart from random flucuations due to natural cell death/birth of cells) until an antigen is injected. In this case the virus SARS-CoV-2 is applied at day 0. It follows a sequence of stochastic events promoting cells duplication, cytokine secretion and eventually culminating in the humoral and cellular immune response. Due to the high degree of details of the angorithms enacting such events, an agent-based model is not described by means of mathematical formulas but rather by *Rules* expressed in natural language without sacrifycing rigor. The Rules of the automaton accounting for the main part of the C-IMMSIM ABM of the immune response to SARS-CoV-2 are listed below. Each rule corresponds to a more-or-less complex algorithm whose details are here neglected because less relevant to the purpose of the present article (more can be found in previous publications of the model).

1. <u>Infection</u>: An infection dose *V*(0)=*V*_0_ is injected into the simulated volume 0
2. <u>Endocytosis</u>: the virus enters epithelial cells (EP)
3. <u>Biosynthesis</u>: the viral RNA and viral proteins are made and assembled into new virions that are released by budding (exocytosis) from infected cells (SARS-CoV-2 follows a *lysogenic cycle*, that is, it does not kill the host). At this stage, infected/injured EP
  - <u>DAMPs release</u>: release danger signal (D) (generally indicating interferon, cytokines, DAMPs = damage associated molecular patterns)
  - <u>Inflammation</u>: release IL-6
  - <u>Endocytic presentation</u>: process the viral proteins leading to their presentation on class I HLA molecules
4. <u>B phagocytosis</u>: B cells phagocyte, internalise, process and present viral peptides on class II HLA
5. Response to Danger:
  - <u>NK response</u>: Natural killer cells (NKs) release IFNg upon bystander stimulation by danger
  - <u>M response</u>: Macrophages (M) respond to danger (e.g., DAMPs) via TLR4 releasing TNFa and IL-6
6. <u>M activation</u>: macrophages become activated by IFNg (activated M have a greater phagocytic activity)
7. Active M
  - <u>M phagocytosis</u>: M internalise, process and present viral peptides on class II HLA; in presence of IFNg they release IL-12; they also release TNFa
  - <u>DC activation</u>: M release TNFa which activate dendritic cells (DC)
8. <u>DC phagocytosis & endocytosys</u>: DC phagocyte, internalise, process and present viral peptides on class II HLA (exocytic pathway) but also on class I HLA (endocytic pathway)
9. <u>Th activation</u>: in presence of danger signal, resting T helper lymphocytes are activated by interaction with peptide-bound HLAs on professional antigen presenting cells (M and DC, mainly DC) surface by means of specific interaction with their T-cell receptors (TCR); if no danger is present, the Th cells becomes anergic upon interaction of its TCR with the HLApepide complex
10. <u>Th stimulation by APCs</u>: activated Th interacting with antigen presenting cells (M, DC)
  - <u>Th duplication</u>: start clone expansion; part of the clones become memory cells
  - Th cells release IL-2
  - M release IL-6
  - Th1 release IFNg
  - Th2 release IL-4
  - release IL-12 in presence of high local concentration of IFNg
  - Treg release TGFb and IL-10
11. <u>Th stimulation by B</u>: activated Th interacting with B cells
  - <u>B duplication</u>: stimulate B cells to start clone expansion; part of the clone become memory
  - <u>Th duplication</u>: start clone expansion; part of the clones become memory cells
  - release IL-2, IL-12
  - Th1 release IFNg
  - Th2 release IL-4
  - Treg release TGFb and IL-10
12. <u>Th differentiation</u>: depending on the local concentration of IFNg, IL-10, IL-4, IL-6, IFNb, IL-12, IL-18, IL-2, TGFb and IL23, active T helper cells undergo class switch into Th1 and Th2
13. <u>B differentiation</u>: B cells differentiate to antibody-secreting plasma B cells (PLB)
14. <u>Isotype switch</u>: B cells perform immunoglobulin class switching, that is, change production of immunoglobulin from the isotype IgM to the isotype IgG
15. <u>Antibodies production</u>: Plasma cells secrete antibodies
16. <u>Humoral response</u>: antibodies inhibit viral particles by opsonization; the result are the immuno-complexes that are eventually cleared by macrophages
17. <u>Tc activation</u>: in presence of IL-2, resting cytotoxic T cells (Tc) are activated by the interaction of their TCR with DC presenting on class I HLA the viral peptides but only in presence of IL-2
18. <u>Tc duplication</u>: activated Tc interact with infected EP cells presenting viral peptides on class I HLA molecule
  - <u>Cytotoxic response</u>: kill infected EP (this will further release danger signal)
  - Tc start duplication

## Appendix B. HLA class I peptide list

Each column reports the peptides relative to the allele indicated in the first row. Each entry of the table shows the peptide, the rank score and the relative amino acid position. The relative rank score is used to directly compute the probability to successful bind the peptide to the HLA molecule thus presenting the HLA-peptide complex to the cell surface.

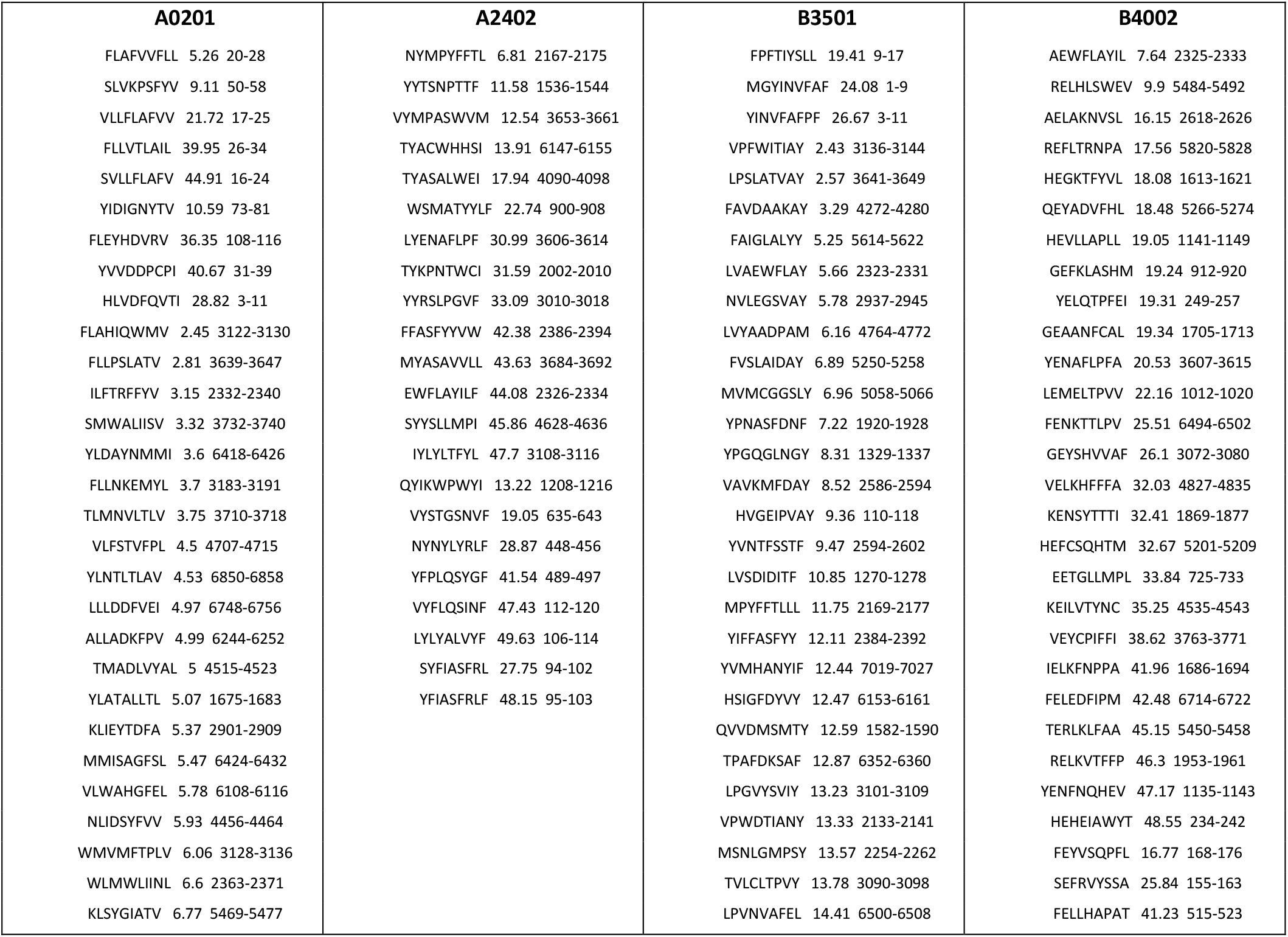
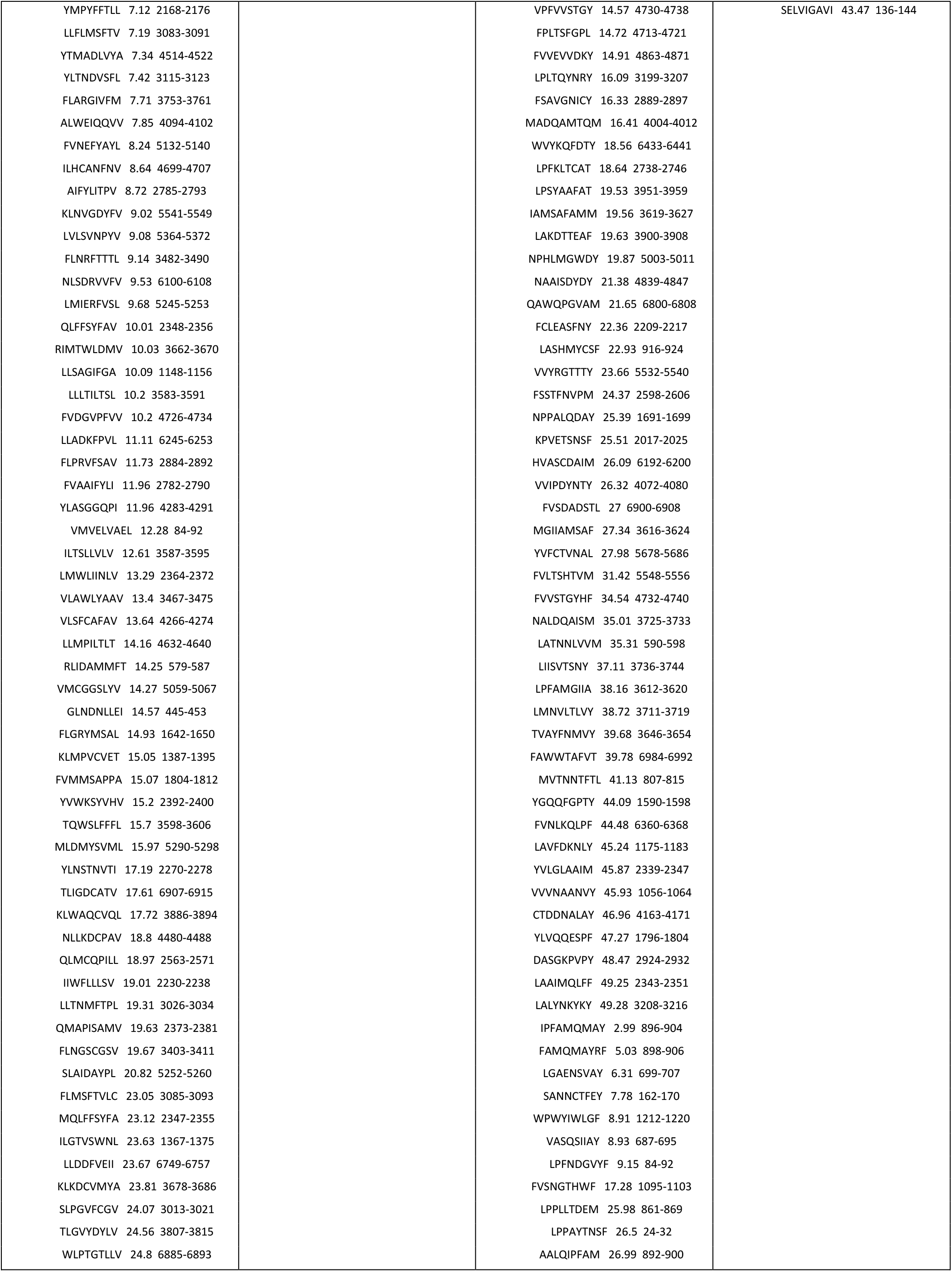
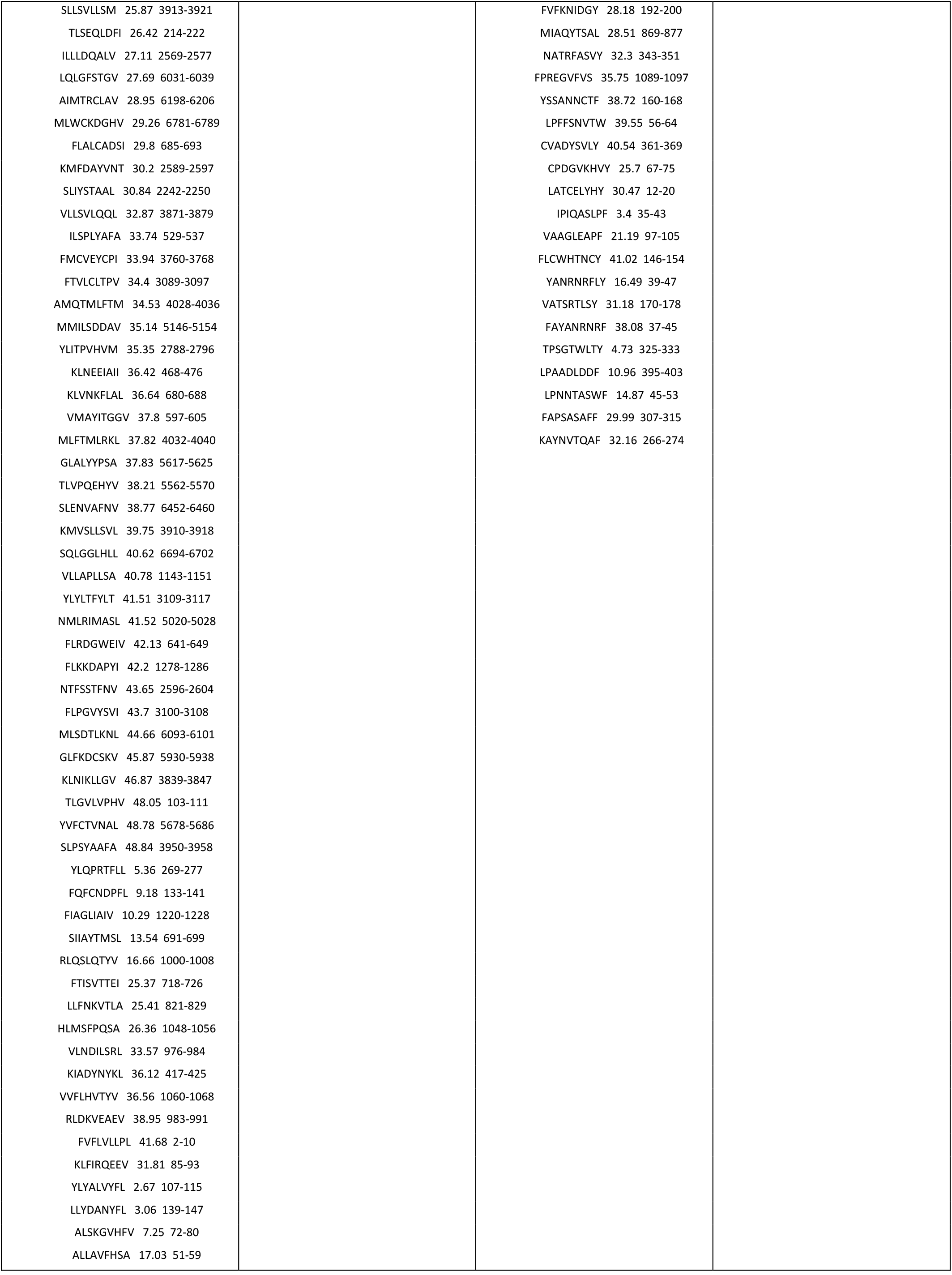
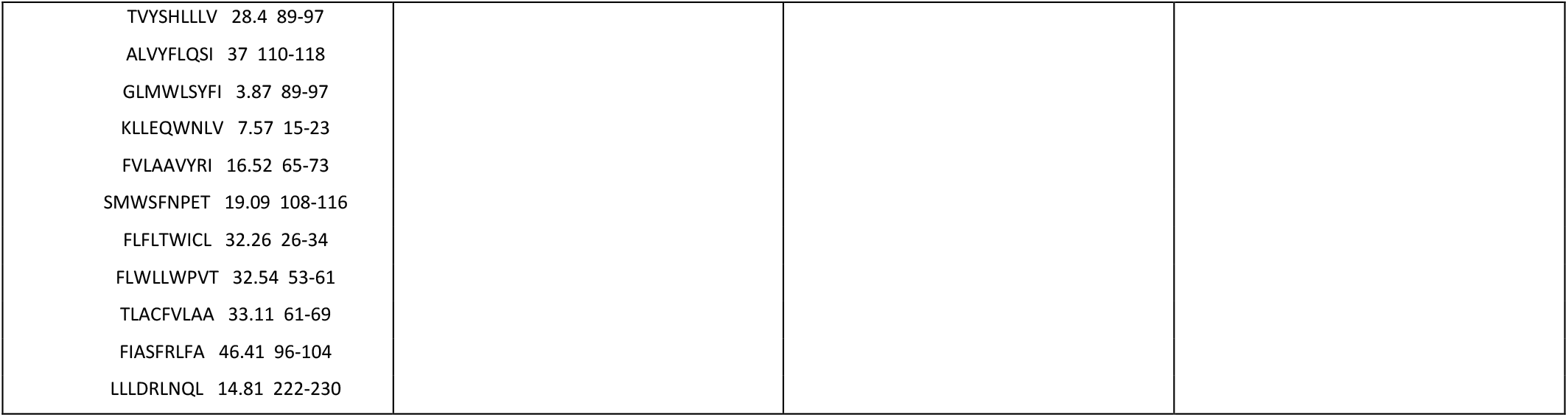

## Appendix C. HLA class II peptides

Each column reports the peptides relative to the HLA indicated in the first row. Each entry of the table shows the peptide and the relative rank score. The relative rank score is used to directly compute the probability to successful bind the peptide to the HLA molecule thus presenting the HLA-peptide complex to the cell surface.

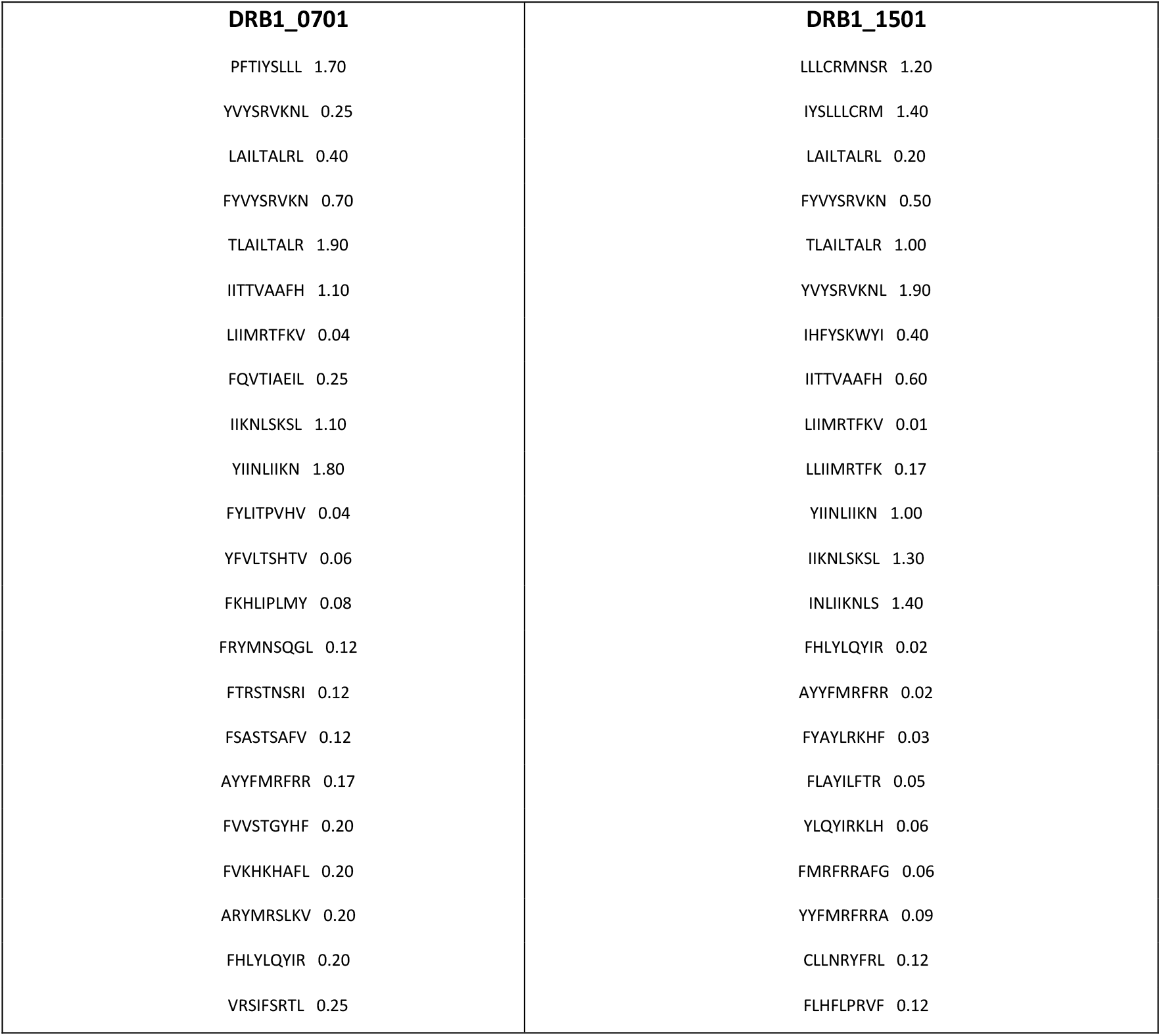
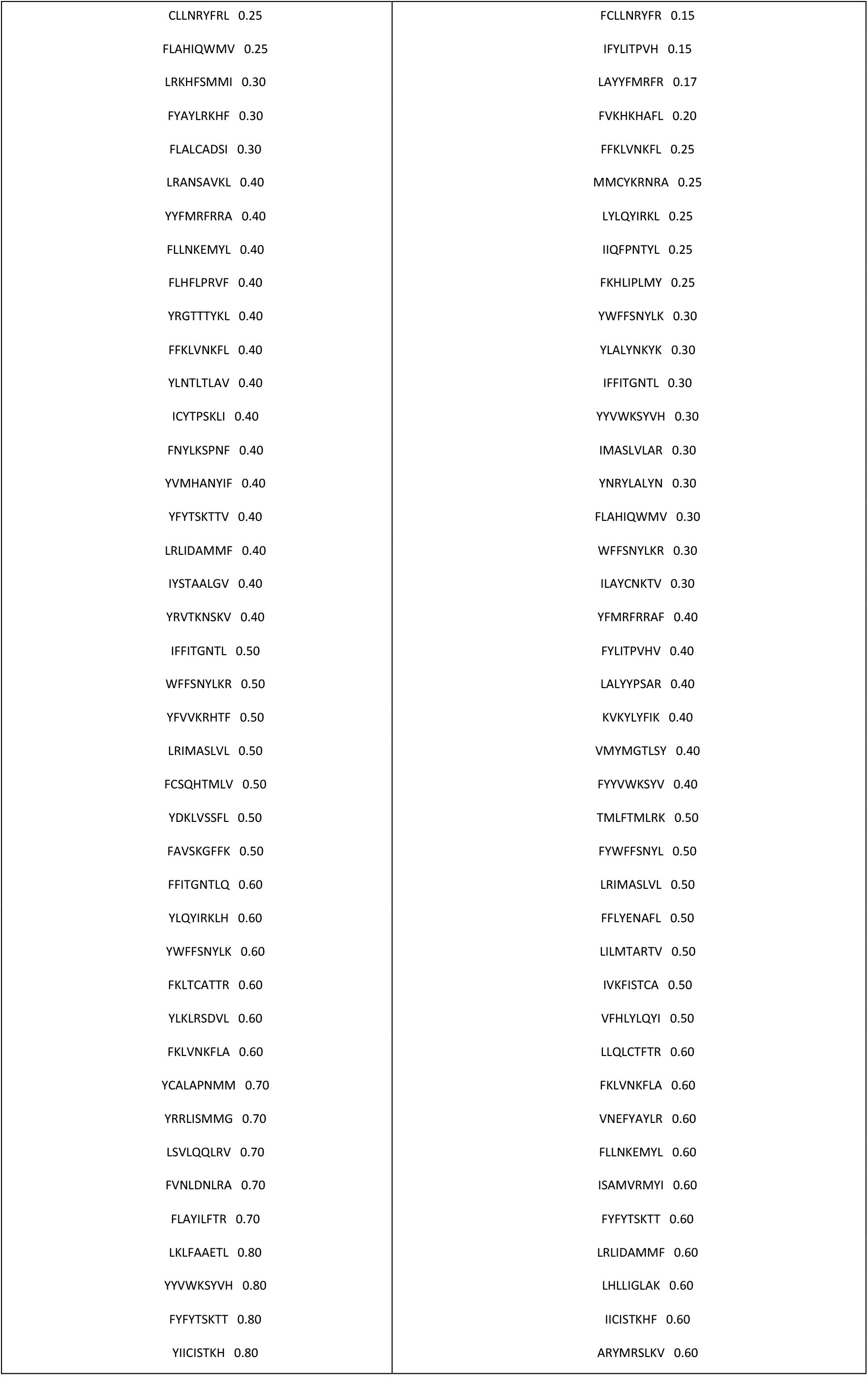
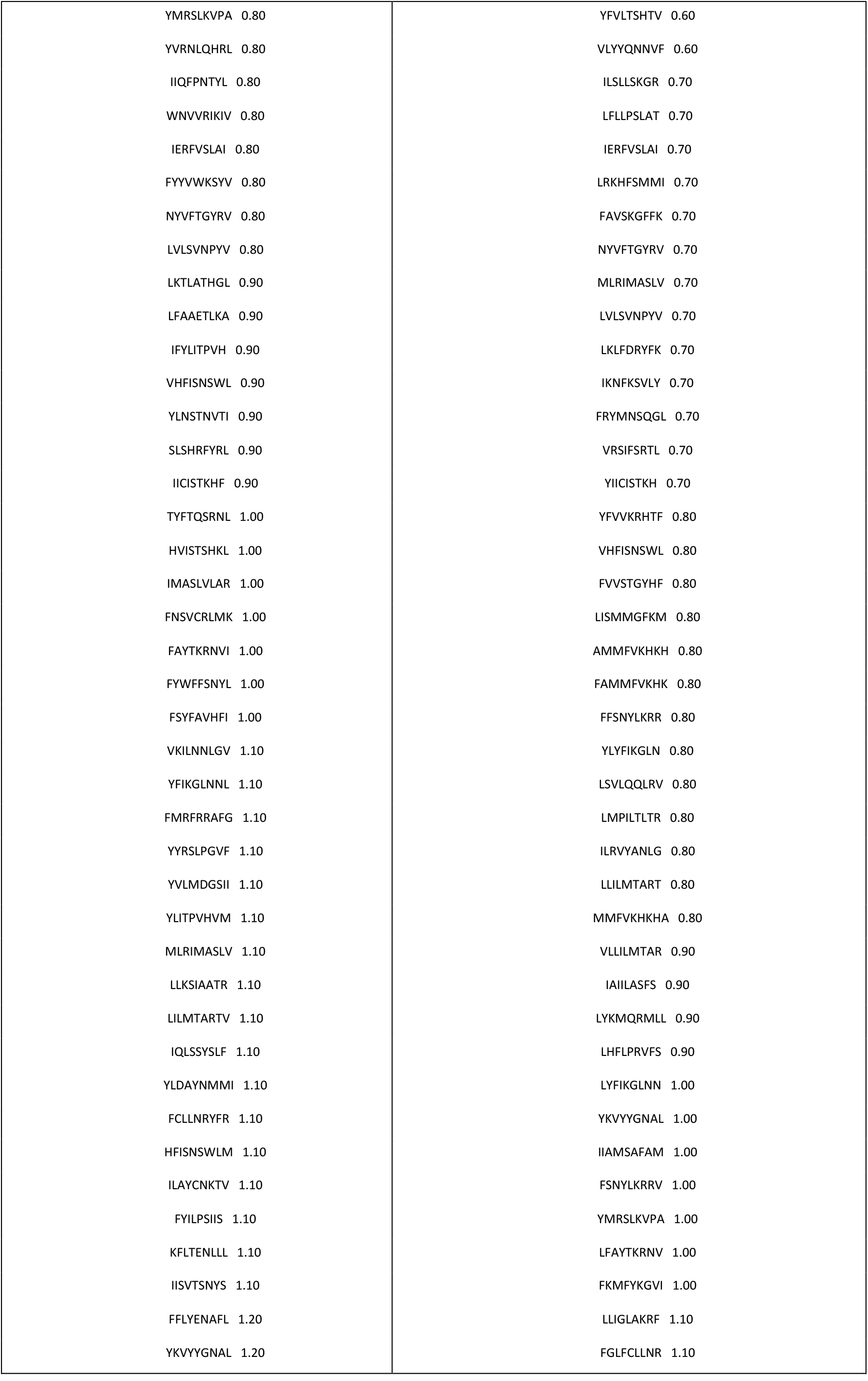
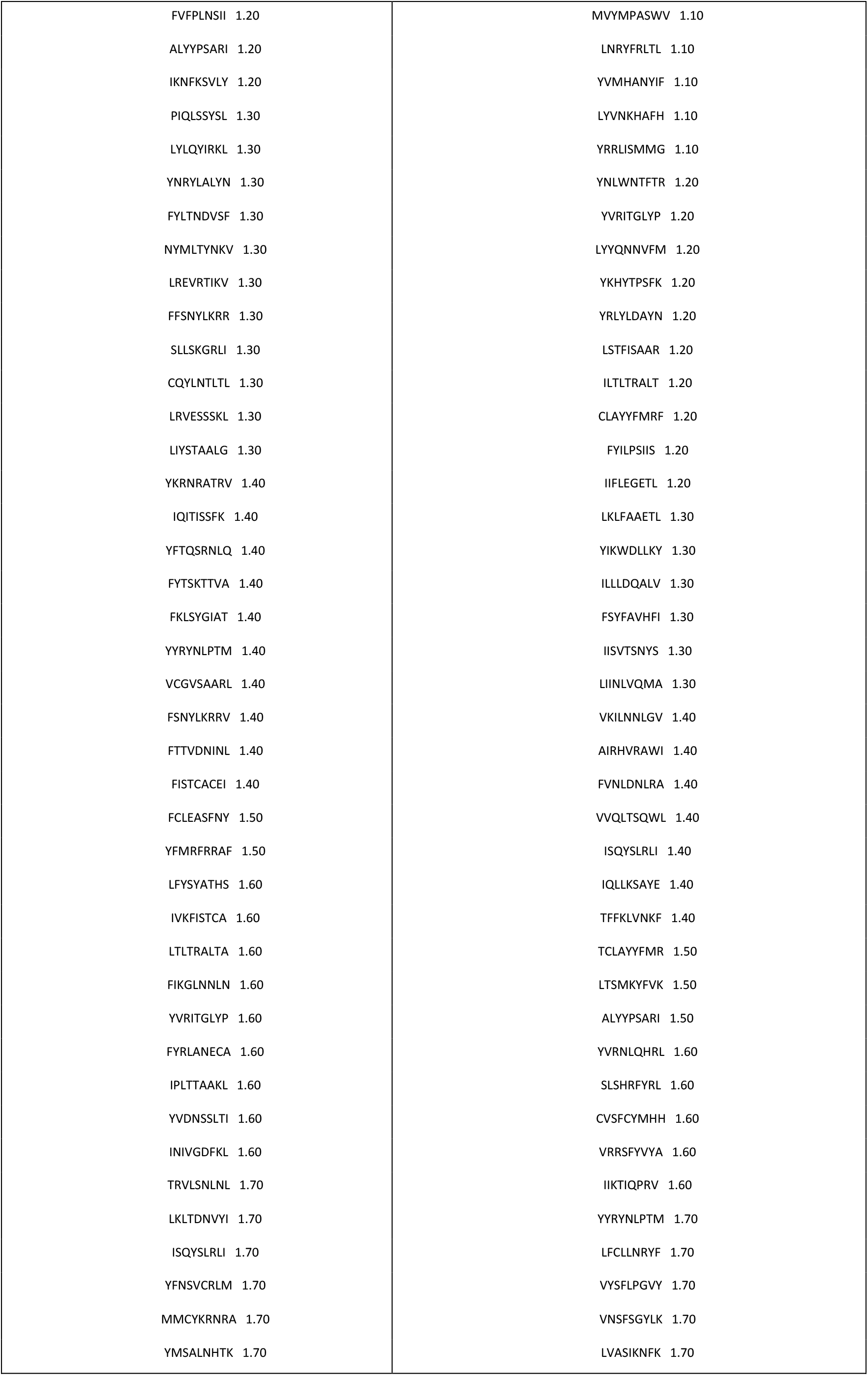
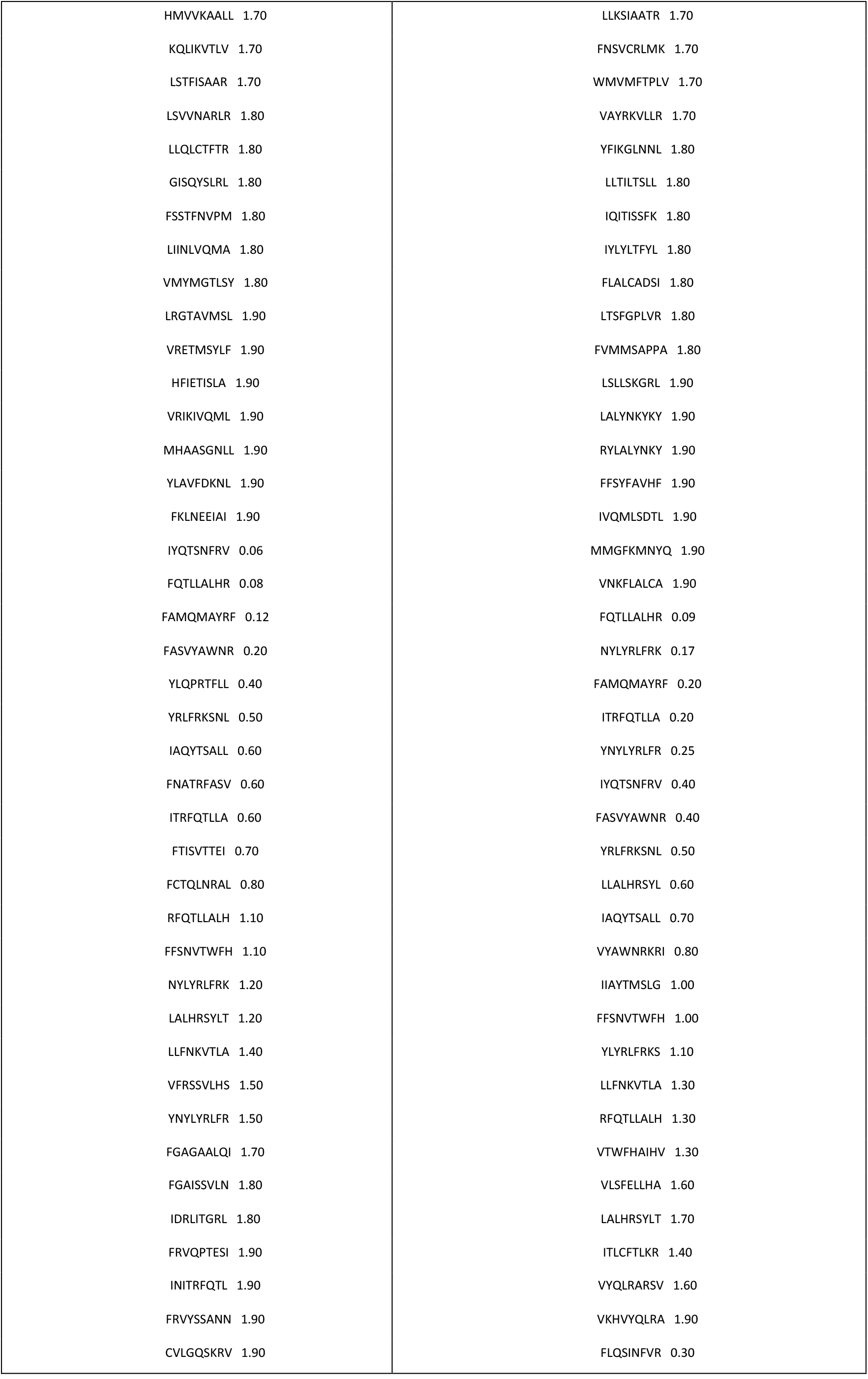
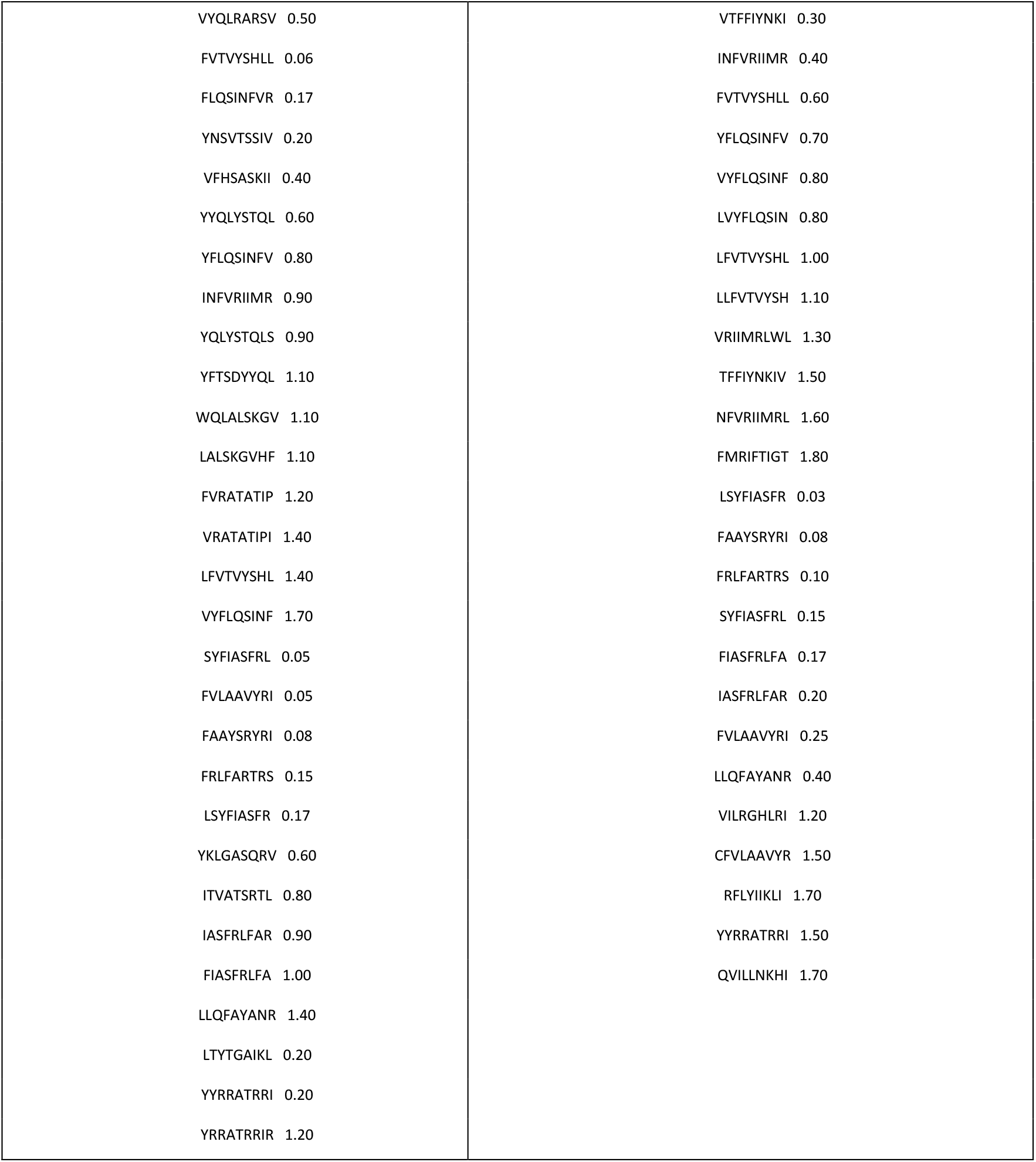

## Appendix D. Example readout of a simulation

Figure 15 shows in panel A how the viral load *V*(*t*) (both soluble, meaning outside infected cells, and proviral, meaning inside infected cells, and the sum of the two), varies with time: the viral particles of SARS-Cov-2 injected at day 0, peaking at about day 5 and start to decline after that and in correspondance to the appearance of antibody producing plasma cells (panel B of Figure 16). In the same panel the immunoglobulins titers are plotted (split in IgM and IgG, further split in IgG1 and IgG2) together with the immunocomplexes (IC) that are antibodies bound to viruses (i.e., opsonised viruses). This plots shows a humoral response surging at abouty day 12 and clearing the virus in about five days. It also shows that the antibody levels remains high for some time after the simulation ends at day 30.

**Figure 15.**
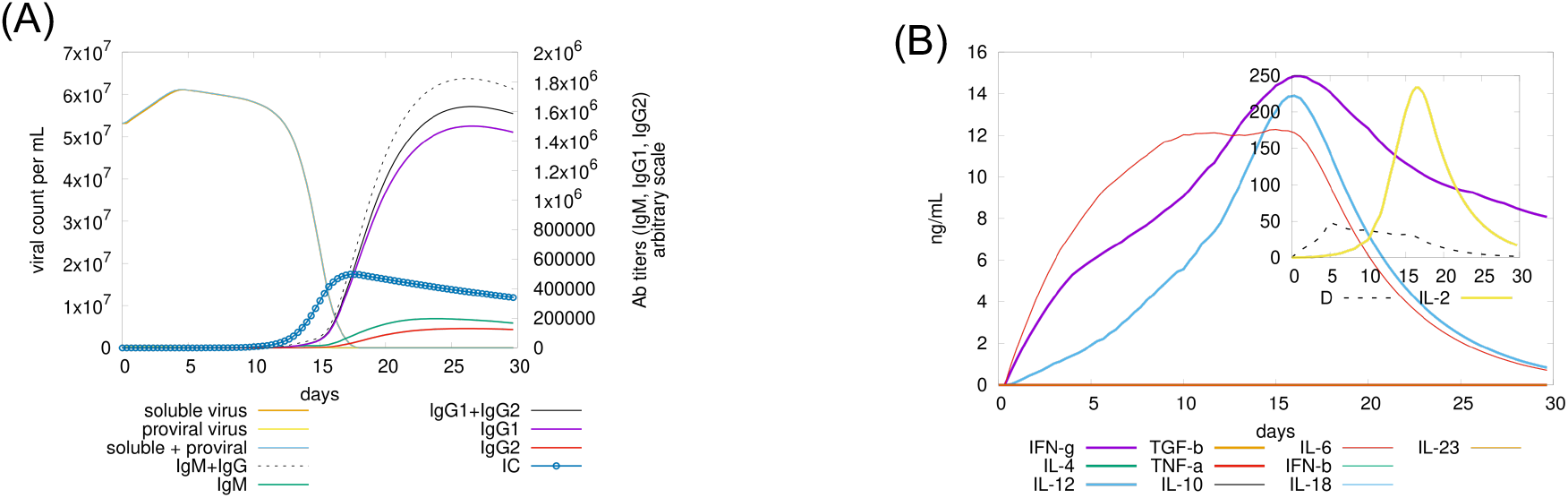
Viral load *V*(*t*), immunoglobulins (IgM and IgG) and the immunocomplexes (IC) are shown in panel A. The same panel shows the viral particles of SARS-Cov-2 injected at day 0, peaking at about day 5 and start to decline after that and in correspondance to the appearance of antibody producing plasma cells (panel B of Figure 16). Panel B shows the cytokines generated during the immune response (see text for details).

**Figure 16.**
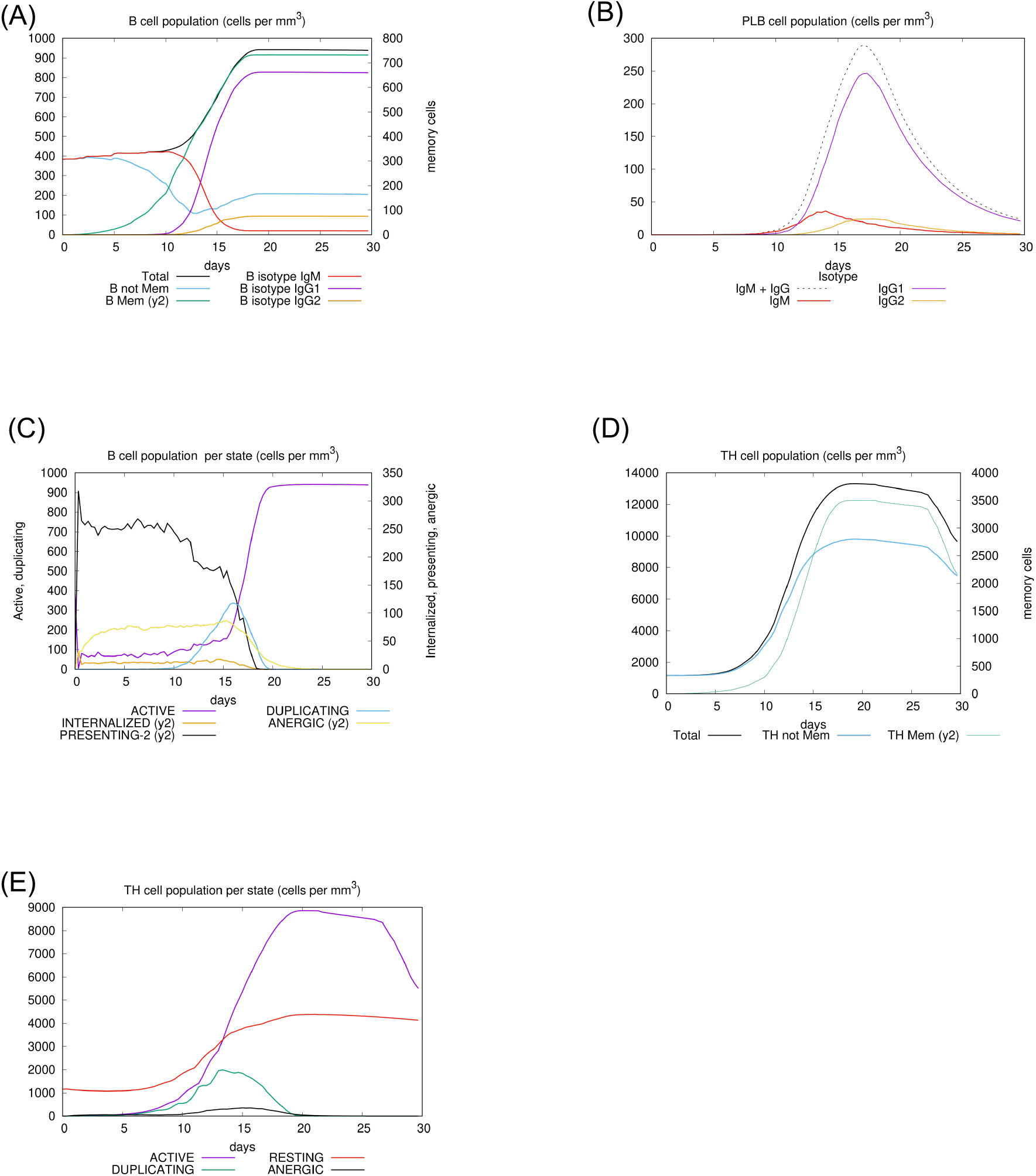
This figure shows details of the population dynamics of B-cells (panel A and C), plasma B cells (PLB in panel B) and helper T lymphocytes (panel D and E).

Panel (B) plots in arbitrary scale the cytokine concentrations for interleukins, interferon and danger signal (inset plot of the same panel). In particular this run shows a high level of IFNg, IL-6, IL-2 and IL-12 elicited by the infection besides a moderate production of danger signal (D, in the inset plot of panel B). Release of cytokines follows from the dynamical rules characterising the agent’s behaviour reported in Appendix A.

Panel A of Figure 16 shows the total counts of the B-cells in all phenotypes, that is, memory and not memory, and the three isotypes IgM, IgG1 and IgG2 (i.e., cells that will become plasma B cells producing IgM or IgG antibodies).

Panel B shows the count of antibody-generating plasma cells subdivided in the three classes IgM, IgG1 and IgG2.

Panel C gets into the details of the ABM simulation model by showing the counts of B-cells subdivided according to the internal state assemed. Worth to note that the cells enter the duplication state only after day 10 and until day 20 because besides presenting the viral peptides on their HLA molecules, they need to be stimulated by stimulated cognate helper T cells bearing the “correct” cell receptor (panel D of same Figure 16). Note also that upon clearance of the virus (cf. panel A of Figure 15) the B-cell population switches back to the “active” state and terminates the presentation of the viral proteins.

Panel D shows cell counts for CD4 T-cell population (total number, memory cells, not memory). Some days after the infection and upon successful interaction with antigen presenting cells, T helper lymphocytes start to duplicate and differentiate into memory. They also foster cytotoxicity (cf. panel A and B of Figure 17) and humoral response (cf. panel B of Figure 16) through secretion of cytokines (panel B of Figure 15). As for B cells in panel A, panel E shows Th counts specifying the internal state of the lymphocyte thus revealing the “activation” phase starting quite soon before day 5 and the duplication phase starting immediately after and ending at about day 20. Of note, some cell become anergic around day 15 for lack of “danger” (second) signal upon activation (Rule n.9 in Appendix A).

**Figure 17.**
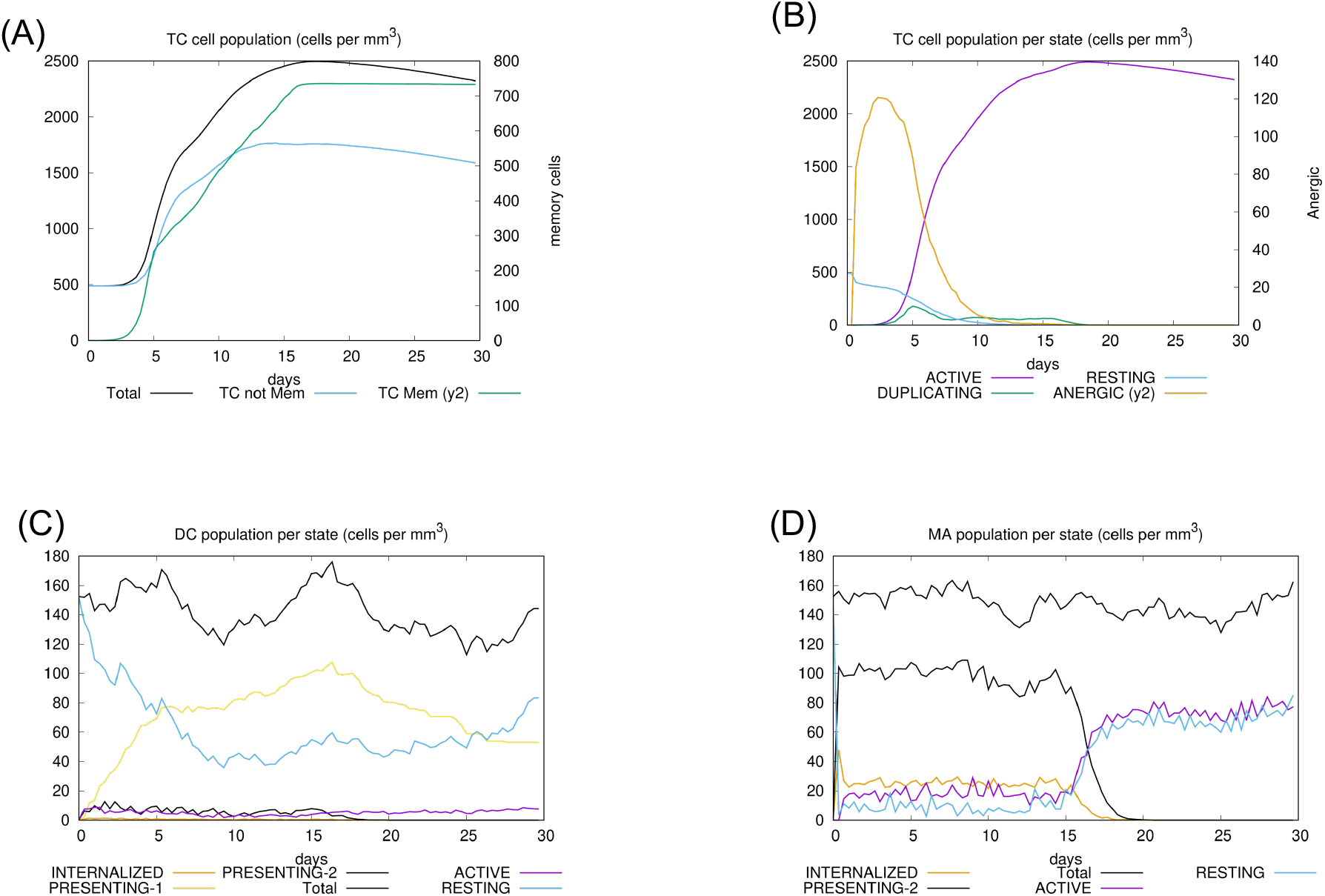
Here we plot the cell counts and detailed intra-cellular state numbers of cytotoxic T cells (panel A and B) and of antigen presenting cells DC and M (respectively in panels C and D). Further details in the text.

Panel A of Figure 17 plots the cell counts for CD8 T-cell population (total number, memory cells, not memory). Panel B shows the counts per internal cell state revealing a limited number of anergic cells (numbers on the y2-axis), the early activation of a large number from day 3 (and corresponding decrease of resting cells) entering a duplication phase peaking ad day 5 but progressing through about day 18.

Panels C and D show the total number and the breakout counts for antigen presenting cells DC (denritic cells in panel C) and MA (that we also indicated as M, macrophages, in panel D). In particular the presentation activity following the internalization of the virus by macrophages terminates at about day 20. Similar behaviour is shown in panel E for dendritic cells. Interestingly more macrophages are activated following Rule n.6 (in Appendix A) due to the high level of IFNg released by natural killer cells (cf. Rule n.5 in Appendix A) upon bystander stimulation by danger signals (or damage associated molecules, cf. panel B of Figure 15) segreted by infected cells.

## References

[1] F. Castiglione and F. Celada, Immune System Modeling and Simulation. Boca Raton: CRC Press, 2015.

[2] H. Takaba and H. Takayanagi, “The Mechanisms of T Cell Selection in the Thymus,” Trends in Immunology, vol. 38, no. 11, pp. 805–816, Nov. 2017, doi: 10.1016/j.it.2017.07.010.

[3] F. Castiglione, D. Santoni, and N. Rapin, “CTLs’ repertoire shaping in the thymus: a Monte Carlo simulation.,” Autoimmunity, vol. 44, no. 4, pp. 261–70, Jun. 2011, doi: 10.3109/08916934.2011.523272.

[4] M. Burnet, “The Clonal Selection Theory of Acquired Immunity,” The Yale journal of biology and medicine, 1960, doi: 10.1016/j.juro.2012.06.070.

[5] A. M. Silverstein, “The Clonal Selection Theory: what it really is and why modern challenges are misplaced,” Nature Immunology, vol. 3, no. 9, pp. 793–796, Sep. 2002, doi: 10.1038/ni0902-793.

[6] J. Lederberg, “Genes and antibodiesGenes and Antibodies: Do antigens bear instructions for antibody specificity or do they select cell lines that arise by mutation?,” Science, vol. 129, no. 3364, pp. 1649–1653, 1959, doi: 10.1126/science.129.3364.1649.

[7] S. Brenner and C. Milstein, “Origin of antibody variation,” Nature, vol. 211, no. 5046, pp. 242–243, 1966, doi: 10.3109/00365526809180138.

[8] S. Tonegawa, “Somatic generation of antibody diversity,” Nature, vol. 302, no. 5909, pp. 575–581, Apr. 1983, doi: 10.1038/302575a0.

[9] F. N. Papavasiliou and D. G. Schatz, “Somatic Hypermutation of Immunoglobulin Genes,” Cell, vol. 109, no. 2, pp. S35–S44, Apr. 2002, doi: 10.1016/S0092-8674(02)00706-7.

[10] L. Hayflick and P. S. Moorhead, “The serial cultivation of human diploid cell strains,” Experimental Cell Research, vol. 25, no. 3, pp. 585–621, Dec. 1961, doi: 10.1016/0014-4827(61)90192-6.

[11] J. W. Shay and W. E. Wright, “Hayflick, his limit, and cellular ageing,” Nature Reviews Molecular Cell Biology, vol. 1, no. 1, pp. 72–76, Oct. 2000, doi: 10.1038/35036093.

[12] R. H. Schwartz, “T cell anergy,” Annual Review of Immunology, vol. 21, no. 1, pp. 305–334, Apr. 2003, doi: 10.1146/annurev.immunol.21.120601.141110.

[13] S. D. Saibil, E. K. Deenick, and P. S. Ohashi, “The sound of silence: modulating anergy in T lymphocytes,” Current Opinion in Immunology, vol. 19, no. 6, pp. 658–664, Dec. 2007, doi: 10.1016/j.coi.2007.08.005.

[14] G. J. v. Nossal and B. L. Pike, “Clonal anergy: Persistence in tolerant mice of antigen-binding B lymphocytes incapable of responding to antigen or mitogen,” Proceedings of the National Academy of Sciences, vol. 77, no. 3, pp. 1602–1606, Mar. 1980, doi: 10.1073/pnas.77.3.1602.

[15] Y. Yarkoni, A. Getahun, and J. C. Cambier, “Molecular underpinning of B-cell anergy,” Immunological Reviews, vol. 237, no. 1, pp. 249–263, Aug. 2010, doi: 10.1111/j.1600-065X.2010.00936.x.

[16] P. Matzinger, “Tolerance, Danger, and the Extended Family,” Annual Review of Immunology, vol. 12, no. 1, pp. 991–1045, Apr. 1994, doi: 10.1146/annurev.iy.12.040194.005015.

[17] S. Gallucci and P. Matzinger, “Danger signals: SOS to the immune system.,” Current opinion in immunology, vol. 13, no. 1, pp. 114–9, Feb. 2001, [Online]. Available: http://www.ncbi.nlm.nih.gov/pubmed/11154927.

[18] T. Pradeu and E. L. Cooper, “The danger theory: 20 years later,” Frontiers in Immunology, vol. 3, 2012, doi: 10.3389/fimmu.2012.00287.

[19] N. K. Jerne, “Towards a network theory of the immune system.,” Annales d’immunologie, 1974, doi: 10.1002/eji.1830050511.

[20] I. Menshikov et al., “The idiotypic network in the regulation of autoimmunity: Theoretical and experimental studies,” Journal of Theoretical Biology, vol. 375, pp. 32–39, Jun. 2015, doi: 10.1016/j.jtbi.2014.10.003.

[21] F. Castiglione, F. Poccia, G. D’Offizi, and M. Bernaschi, “Mutation, fitness, viral diversity, and predictive markers of disease progression in a computational model of HIV type 1 infection,” AIDS Research and Human Retroviruses, vol. 20, no. 12, 2004.

[22] F. Castiglione, K. Duca, A. Jarrah, R. Laubenbacher, D. Hochberg, and D. Thorley-Lawson, “Simulating Epstein-Barr virus infection with C-ImmSim,” Bioinformatics, vol. 23, no. 11, 2007, doi: 10.1093/bioinformatics/btm044.

[23] F. Pappalardo, P.-L. Lollini, F. Castiglione, and S. Motta, “Modeling and simulation of cancer immunoprevention vaccine,” Bioinformatics, vol. 21, no. 12, 2005, doi: 10.1093/bioinformatics/bti426.

[24] J. von Eichborn, A. L. Woelke, F. Castiglione, and R. Preissner, “VaccImm: Simulating peptide vaccination in cancer therapy,” BMC Bioinformatics, vol. 14, 2013, doi: 10.1186/1471-2105-14-127.

[25] F. Castiglione, V. Sleitser, and Z. Agur, “Analysing Hypersensitivity to Chemotherapy in a Cellular Automata Model of the Immune System,” 2003.

[26] V. Prana, P. Tieri, M. C. Palumbo, E. Mancini, and F. Castiglione, “Modeling the Effect of High Calorie Diet on the Interplay between Adipose Tissue, Inflammation, and Diabetes,” Computational and Mathematical Methods in Medicine, vol. 2019, pp. 1–8, Feb. 2019, doi: 10.1155/2019/7525834.

[27] V. Baldazzi, P. Paci, M. Bernaschi, and F. Castiglione, “Modeling lymphocyte homing and encounters in lymph nodes,” BMC Bioinformatics, vol. 10, 2009, doi: 10.1186/1471-2105-10-387.

[28] F. Castiglione, P. Tieri, A. Palma, and A. S. Jarrah, “Statistical ensemble of gene regulatory networks of macrophage differentiation,” BMC Bioinformatics, vol. 17, 2016, doi: 10.1186/s12859-016-1363-4.

[29] F. Castiglione, D. Ghersi, and F. Celada, “Computer Modeling of Clonal Dominance: Memory-Anti-Naïve and Its Curbing by Attrition,” Frontiers in Immunology, vol. 10, Jul. 2019, doi: 10.3389/fimmu.2019.01513.

[30] A. Madonia et al., “Computational modeling of immune system of the fish for a more effective vaccination in aquaculture,” Bioinformatics, vol. 33, no. 19, 2017, doi: 10.1093/bioinformatics/btx341.

[31] T. Kar et al., “A candidate multi-epitope vaccine against SARS-CoV-2,” Scientific Reports, vol. 10, no. 1, p. 10895, Dec. 2020, doi: 10.1038/s41598-020-67749-1.

[32] K. Abraham Peele, T. Srihansa, S. Krupanidhi, V. S. Ayyagari, and T. C. Venkateswarulu, “Design of multi-epitope vaccine candidate against SARS-CoV-2: a in-silico study,” Journal of Biomolecular Structure and Dynamics, pp. 1–9, Jun. 2020, doi: 10.1080/07391102.2020.1770127.

[33] B. Kohler, R. Puzone, P. E. Seiden, and F. Celada, “A systematic approach to vaccine complexity using an automaton model of the cellular and humoral immune systemI. Viral characteristics and polarized responses,” Vaccine, vol. 19, o. 7–8, pp. 862–876, 2000, doi: 10.1016/S0264-410X(00)00225-5.

[34] N. Rapin, O. Lund, M. Bernaschi, and F. Castiglione, “Computational immunology meets bioinformatics: the use of prediction tools for molecular binding in the simulation of the immune system.,” PLoS ONE, vol. 5, no. 4, p. e9862, Jan. 2010, doi: 10.1371/journal.pone.0009862.

[35] H. H. Lin, G. L. Zhang, S. Tongchusak, E. L. Reinherz, and V. Brusic, “Evaluation of MHC-II peptide binding prediction servers: applications for vaccine research.,” BMC bioinformatics, vol. 9 Suppl 12, p. S22, Jan. 2008, doi: 10.1186/1471-2105-9-S12-S22.

[36] H. H. Lin, S. Ray, S. Tongchusak, E. L. Reinherz, and V. Brusic, “Evaluation of MHC class I peptide binding prediction servers?: Applications for vaccine research,” vol. 13, pp. 1–13, 2008, doi: 10.1186/1471-2172-9-8.

[37] O. Lund et al., “Definition of supertypes for HLA molecules using clustering of specificity matrices,” Immunogenetics, vol. 55, no. 12, pp. 797–810, Mar. 2004, doi: 10.1007/s00251-004-0647-4.

[38] M. Nielsen et al., “Improved prediction of MHC class I and class II epitopes using a novel Gibbs sampling approach,” Bioinformatics, vol. 20, no. 9, pp. 1388–1397, Jun. 2004, doi: 10.1093/bioinformatics/bth100.

[39] M. Nielsen, C. Lundegaard, and O. Lund, “Prediction of MHC class II binding affinity using SMM-align, a novel stabilization matrix alignment method,” BMC Bioinformatics, vol. 8, no. 1, p. 238, Dec. 2007, doi: 10.1186/1471-2105-8-238.

[40] F. F. Gonzalez-Galarza et al., “Allele frequency net database (AFND) 2020 update: gold-standard data classification, open access genotype data and new query tools,” Nucleic Acids Research, Nov. 2019, doi: 10.1093/nar/gkz1029.

[41] M. Nielsen et al., “Reliable prediction of T-cell epitopes using neural networks with novel sequence representations,” pp. 1007–1017, 2003, doi: 10.1110/ps.0239403.view.

[42] C. Lundegaard, K. Lamberth, M. Harndahl, S. Buus, O. Lund, and M. Nielsen, “NetMHC-3.0: accurate web accessible predictions of human, mouse and monkey MHC class I affinities for peptides of length 8–11,” Nucleic Acids Research, vol. 36, o. suppl_2, pp. W509–W512, Jul. 2008, doi: 10.1093/nar/gkn202.

[43] M. Andreatta and M. Nielsen, “Gapped sequence alignment using artificial neural networks: application to the MHC class I system,” Bioinformatics, vol. 32, no. 4, pp. 511–517, Feb. 2016, doi: 10.1093/bioinformatics/btv639.

[44] K. K. Jensen et al., “Improved methods for predicting peptide binding affinity to MHC class II molecules,” Immunology, vol. 154, no. 3, pp. 394–406, Jul. 2018, doi: 10.1111/imm.12889.

[45] C. F. A. de Bourcy, C. J. L. Angel, C. Vollmers, C. L. Dekker, M. M. Davis, and S. R. Quake, “Phylogenetic analysis of the human antibody repertoire reveals quantitative signatures of immune senescence and aging,” Proceedings of the National Academy of Sciences, vol. 114, no. 5, pp. 1105–1110, Jan. 2017, doi: 10.1073/pnas.1617959114.

[46] L. Pangrazzi and B. Weinberger, “T cells, aging and senescence,” Experimental Gerontology, vol. 134, p. 110887, Jun. 2020, doi: 10.1016/j.exger.2020.110887.

[47] A. Aiello et al., “Immunosenescence and Its Hallmarks: How to Oppose Aging Strategically? A Review of Potential Options for Therapeutic Intervention,” Frontiers in Immunology, vol. 10, Sep. 2019, doi: 10.3389/fimmu.2019.02247.

[48] M. Cline and JJ. Hutton, Hematology and oncology, Internal medicine. Boston: Little, Brown, 1983.

[49] Z. Liu, X. Bing, and X. Z. Zhi, “The epidemiological characteristics of an outbreak of 2019 novel coronavirus diseases (COVID-19) in China,” Chinese Journal of Epidemiology, vol. 41, no. 2, pp. 145–151, 2020.

[50] G. Onder, G. Rezza, and S. Brusaferro, “Case-Fatality Rate and Characteristics of Patients Dying in Relation to COVID-19 in Italy,” JAMA, Mar. 2020, doi: 10.1001/jama.2020.4683.

[51] Y. Wei et al., “Clinical characteristics of 276 hospitalized patients with coronavirus disease 2019 in Zengdu District, Hubei Province: a single-center descriptive study,” BMC Infectious Diseases, vol. 20, no. 1, p. 549, Dec. 2020, doi: 10.1186/s12879-020-05252-8.

[52] D. Zhao et al., “Asymptomatic infection by SARS-CoV-2 in healthcare workers: A study in a large teaching hospital in Wuhan, China,” International Journal of Infectious Diseases, vol. 99, pp. 219–225, Oct. 2020, doi: 10.1016/j.ijid.2020.07.082.

[53] S.-Y. Zhang et al., “Clinical characteristics of different subtypes and risk factors for the severity of illness in patients with COVID-19 in Zhejiang, China,” Infectious Diseases of Poverty, vol. 9, no. 1, p. 85, Dec. 2020, doi: 10.1186/s40249-020-00710-6.

[54] R. Verity et al., “Estimates of the severity of coronavirus disease 2019: a model-based analysis,” The Lancet Infectious Diseases, vol. 20, no. 6, pp. 669–677, Jun. 2020, doi: 10.1016/S1473-3099(20)30243-7.

[55] M. Cascella, M. Rajnik, A. Cuomo, S. C. Dulebohn, and R. di Napoli, Features, Evaluation, and Treatment of Coronavirus (COVID-19), [Internet]. Treasure Island (FL): StatPearls Publishing.

[56] W. J. Wiersinga, A. Rhodes, A. C. Cheng, S. J. Peacock, and H. C. Prescott, “Pathophysiology, Transmission, Diagnosis, and Treatment of Coronavirus Disease 2019 (COVID-19),” JAMA, vol. 324, no. 8, p. 782, Aug. 2020, doi: 10.1001/jama.2020.12839.

[57] L. F. Westblade et al., “SARS-CoV-2 Viral Load Predicts Mortality in Patients with and without Cancer Who Are Hospitalized with COVID-19,” Cancer Cell, vol. 38, no. 5, Nov. 2020, doi: 10.1016/j.ccell.2020.09.007.

[58] C. M. Petrilli et al., “Factors associated with hospital admission and critical illness among 5279 people with coronavirus disease 2019 in New York City: prospective cohort study,” BMJ, vol. 369, no. BMJ 2020;369:m1966, pp. 1–15, May 2020, doi: 10.1136/bmj.m1966.

[59] X. Chen et al., “Detectable Serum Severe Acute Respiratory Syndrome Coronavirus 2 Viral Load (RNAemia) Is Closely Correlated With Drastically Elevated Interleukin 6 Level in Critically Ill Patients With Coronavirus Disease 2019,” Clinical Infectious Diseases, Apr. 2020, doi: 10.1093/cid/ciaa449.

[60] S. F. Pedersen and Y.-C. Ho, “SARS-CoV-2: a storm is raging,” Journal of Clinical Investigation, vol. 130, no. 5, pp. 2202–2205, Apr. 2020, doi: 10.1172/JCI137647.

[61] C. Lucas et al., “Longitudinal analyses reveal immunological misfiring in severe COVID-19,” Nature, vol. 584, no. 7821, pp. 463–469, Aug. 2020, doi: 10.1038/s41586-020-2588-y.

[62] A. L. Mueller, M. S. McNamara, and D. A. Sinclair, “Why does COVID-19 disproportionately affect older people?,” Aging, vol. 12, no. 10, May 2020, doi: 10.18632/aging.103344.

[63] G. Chen et al., “Clinical and immunological features of severe and moderate coronavirus disease 2019,” Journal of Clinical Investigation, vol. 130, no. 5, pp. 2620–2629, Apr. 2020, doi: 10.1172/JCI137244.

[64] K. L. Lynch et al., “Magnitude and Kinetics of Anti–Severe Acute Respiratory Syndrome Coronavirus 2 Antibody Responses and Their Relationship to Disease Severity,” Clinical Infectious Diseases, Jul. 2020, doi: 10.1093/cid/ciaa979.

[65] M. Lisboa Bastos et al., “Diagnostic accuracy of serological tests for covid-19: systematic review and meta-analysis,” BMJ, p. m2516, Jul. 2020, doi: 10.1136/bmj.m2516.

[66] F. Muecksch, H. Wise, and et al., “Longitudinal analysis of clinical serology assay performance and neutralising antibody levels in COVID19 convalescents,” medRxiv, vol. Aug 6, 2020.

[67] L. R. Dice, “Measures of the Amount of Ecologic Association Between Species,” Ecology, vol. 26, no. 3, pp. 297–302, Jul. 1945, doi: 10.2307/1932409.

